# Discovery and heterologous expression of functional 4-O-dimethylallyl-L-tyrosine synthases from lichen-forming fungi

**DOI:** 10.1101/2024.03.23.586307

**Authors:** Riccardo Iacovelli, Siqi He, Nika Sokolova, Iris Lokhorst, Maikel Borg, Peter Fodran, Kristina Haslinger

**Author notes:** **Corresponding Author Kristina Haslinger –** Department of Chemical and Pharmaceutical Biology, Groningen Research Institute of Pharmacy, University of Groningen, 9713 AV Groningen, The Netherlands. These authors contributed equally (Siqi He, Nika Sokolova).

## Abstract

Fungal DMATS-type aromatic prenyltransferases are a family of biosynthetic enzymes that catalyze the prenylation of a range of aromatic substrates during the biosynthesis of bioactive indole alkaloids, diketopiperazines, and meroterpenoids. Together with their broad substrate scope and soluble nature, this makes DMATS-type prenyltransferases particularly adept for applications in biocatalysis, for example to derivatize aromatic drug leads and improve their bioactivity. Here, we investigated four putative DMATS-type prenyltransferases from lichen-forming fungi, an underexplored group of organisms that produce more than 1,000 unique metabolites. We were able to successfully express two functional lichen prenyltransferases in the heterologous host *A. oryzae*, which allowed us to identify them as 4-O-dimethylallyltyrosine synthases. Our extensive bioinformatic analysis shows that related lichen prenyltransferases are likely not active on indoles but rather on aromatic polyketides and phenylpropanoids, common metabolites in these organisms. Overall, our work not only provides new insights into fungal DMATS-type prenyltransferases at the family level, but it also enables future efforts aimed at identifying new candidates for biocatalytic transformations of aromatic compounds.

## INTRODUCTION

Fungi are proficient producers of secondary metabolites: specialized molecules that are not strictly required for growth or survival, but that play important roles in sexual development, communication and competition with other organisms, and defense from environmental stressors [1]. In particular, lichen-forming fungi—organisms living in symbiotic associations with algae or cyanobacteria in nature—are prolific producers of secondary metabolites, notably bioactive aromatic polyketides and natural dyes [2–5]. Lichens are relatively underexplored because of their complex biology, as well as the technical challenges around cultivation in laboratory settings [6]. Thanks to advancements in DNA sequencing technologies and bioinformatics, it has recently become easier to obtain high-quality genome sequences from lichen-forming fungi and thus gauge their biosynthetic capabilities [7–9]. A major challenge, however, remains the heterologous expression of genes originating from these organisms, thus hampering the linking of metabolites to biosynthetic genes [10,11]. To date, there are only three studies reporting the successful expression of lichen genes in heterologous hosts: the *pyrG* decarboxylase from *Solorina crocea* [12]; and two polyketide synthase-encoding genes from *Pseudevernia furfuracea* [13]and *Stereocaulon alpinum* [14]. Despite this challenge, we set out to explore the biosynthetic potential of lichen-forming fungi experimentally and thus analyzed their predicted biosynthetic gene clusters *in silico*. Surprisingly, we found that 67 genes were annotated as dimethylallyltryptophan synthase (DMATS), biosynthetic enzymes belonging to the superfamily of ABBA-type prenyltransferases (PTs) [15–17]. DMATSs are known to be involved in the biosynthesis of indole alkaloids—compounds with potent biological activities in animals, most notably on the nervous and circulatory systems [18]. However, to the best of our knowledge, no indole alkaloids have ever been reported in extracts of lichens [19].

Intrigued by this finding, we selected 4 putative DMATS-type PTs from lichen-forming fungi and overexpressed them in the heterologous host *A. oryzae* for *in vivo* characterization. We furthermore carried out an extensive bioinformatic analysis to investigate sequence-function relationships in this important family of enzymes. The combined results revealed that the prenyltransferases from lichen-forming fungi have a distinct substrate scope that sets them apart from “prototype” tryptophan DMATS, with potential implications in the discovery of new biocatalysts and new bioactive compounds.

## RESULTS AND DISCUSSION

Since we were already working with the wolf lichen *Letharia lupina* in the context of another study [20], we analyzed the biosynthetic gene clusters (BGCs) in its recently published genome [7] with the antiSMASH webtool [21,22]. Among the expected high number of polyketide-producing BGCs (21 out of 42, in total), we also noticed 2 gene clusters annotated as indole-producing, with DMATS as the core biosynthetic enzyme. Intrigued by this finding, we analyzed all published lichen genomes for the presence of genes annotated as putative DMATS and found 67 genes. When further analyzing the genomic context of these genes by antiSMASH and gggenomes [23], two genomic regions struck us as interesting: region AS-522.1 in the genome of *Acarospora strigata* CBS132363, annotated to encode a di-domain DMATS-cytochrome P450 enzyme; region RI-146.1 in the genome of *Ramalina intermedia* YAF0013, annotated to encode a di-domain halogenase-acyltransferase enzyme in the vicinity of the DMATS-encoding gene (Table S1, Figure S1). In addition to these two intriguing enzymes, we also chose the putative DMATS-encoding genes KAF6225851.1of *L. lupina* and its close relative KAF6239039.1 from *L. columbiana* for further characterization by heterologous expression in *Aspergillus oryzae* NSAR1 [24]. When cloning the target gene from the genomic DNA of *A. strigata*, we noticed that all clones were missing 28 nucleotides in the predicted exon 2, which results in a frameshift shortening the open reading frame to 1,358 nucleotides. This is the expected size for a DMATS-encoding gene, and we therefore concluded that the gene was in fact not coding for a di-domain enzyme. We proceeded with the expression of the four DMATS coding sequences.

### Heterologous expression of two lichen DMATS in *A. oryzae* yields 4-O-dimethylallyl-tyrosine and its N-acetyl derivative

We first obtained the synthetic gene encoding *Ri* DMATS, while the sequences encoding *As* DMATS, *Ll* DMATS and *Lc* DMATS were amplified directly from genomic DNA of *A*. strigata CBS 132363 and wild isolates of *L. lupina* and *L. columbiana* collected for a previous study [20]. All genes of interest were cloned into the pTYargB or pTYadeA vectors [25] for overexpression in the host *Aspergillus oryzae* NSAR1. The transformed strains were first grown for 5 days on inducing medium, followed by extraction of secondary metabolites from both mycelium and agar. HPLC-MS-DAD analysis did not reveal the production of any new metabolites in the *Ll* DMATS and *Lc* DMATS overexpression strains compared to the control. Conversely, both *Ri* DMATS and *As* DMATS overexpression led to the production of two new compounds (**1** and **2**) with m/z 250 and 292, respectively (Fig. 1a), albeit at much different levels. Indeed, the concentration of **1** and **2** in the extracts from the *Ri* DMATS overexpression strain are ∼30 higher (Fig. 1b). As neither m/z values matched the expected value for dimethylallyltryptophan (m/z = 273 [M + H]^+^), we subjected the same samples to HRMS-MS^2^ to attempt compound identification. Compound **1** was predicted to have the molecular formula C_14_H_19_NO_3_ based on m/z 250.1438 [M + H]^+^ (calculated for [M + H]^+^, 250.1443), consistent with the attachment of a dimethylallyl moiety to the amino acid tyrosine. Indeed, MS^2^ data showed the typical fragmentation pattern of tyrosine (Table S2, Fig. S8). Compound **2** was predicted to have the molecular formula C_16_H_21_NO_4_ based on m/z 292.1547 [M + H]^+^ (calculated for [M + H]^+^, 292.1549), consistent with the attachment of a dimethylallyl moiety to an acetylated derivative of tyrosine. Once again, MS^2^ data confirmed the hypothesis showing the fragmentation pattern of N-acetyl-tyrosine, a naturally occurring derivative of the amino acid (Table S2, Fig. S9).

**Figure 1.**
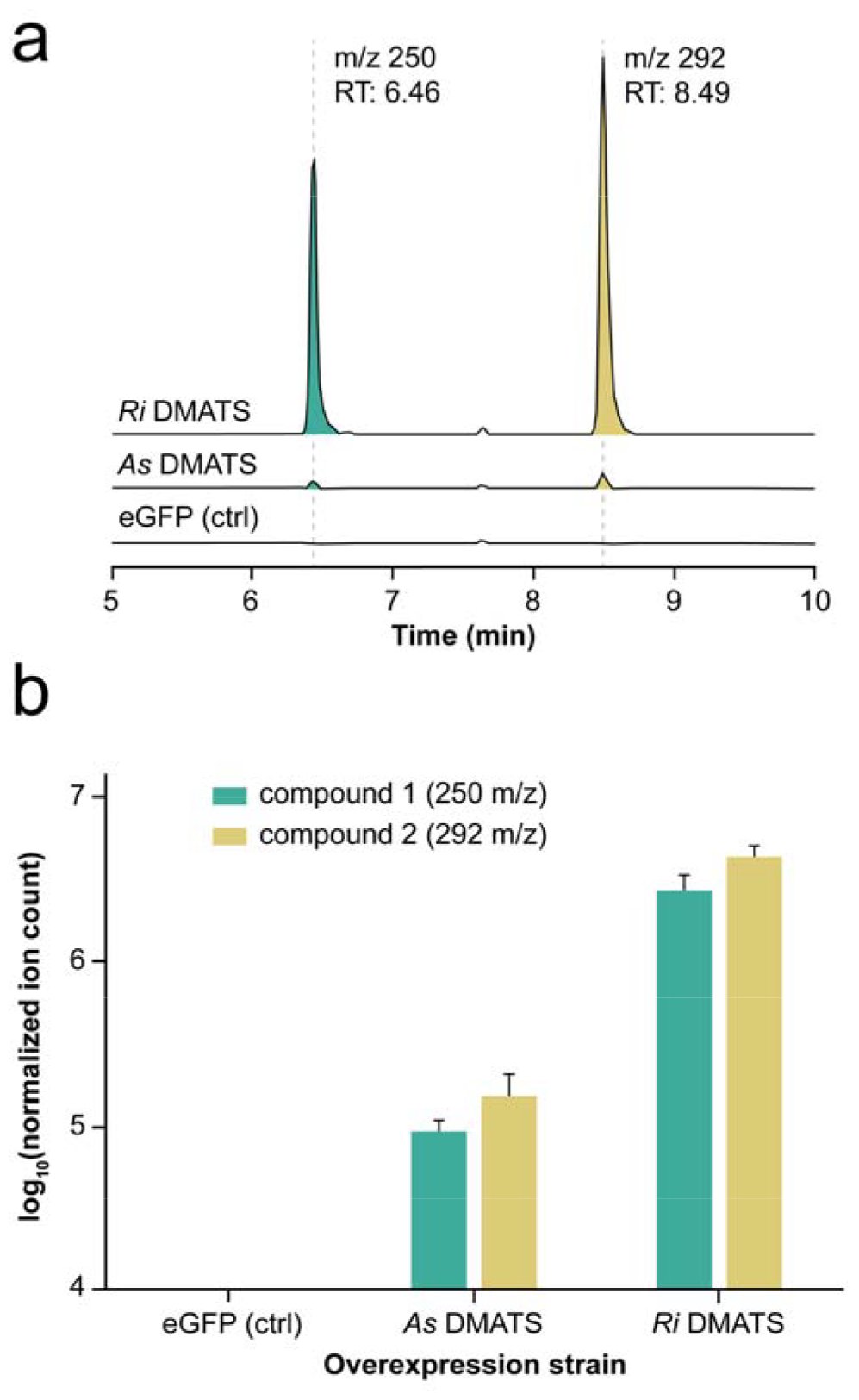
LC-MS–based detection of two new peaks in the extracts of DMATS overexpression strains. **(a)** Chromatograms of fungal extracts displaying EICs of m/z 250 (**1**) and m/z 292 (**2**) (± 0.2 Da). **(b)** Peak areas (log_10_ scale) of **1** and **2** in fungal extracts. The minimum value for the y axis equals a signal intensity of 1 × 10^4^, which corresponds to baseline levels. The values were normalized for the internal standard caffeine. Bars represent mean ± SD (n = 4).

To determine the site of prenylation, we set out to purify **1** and **2** from the total fungal extract of the *Ri* DMATS overexpression strain and perform ^1^H NMR. Because the compounds are only produced on solid media and in limited amounts, we first attempted purification via semi-preparative HPLC to be able to work with smaller culture and solvent volumes. By applying one round of semi-preparative HPLC on an RP-C18 column, we obtained a relatively pure “enriched” fraction containing both compounds **1** and **2** from a total volume of 1 liter of cultivation media and mycelia (Fig. 2a). Given the structural similarity of the two compounds, we hypothesized that their ^1^H NMR spectra would largely overlap, and therefore we submitted the enriched fraction without further purification. The resulting spectrum (Fig. 2b and S2) clearly shows that the aromatic rings of both compounds are free of substitutions, indicating that the site of prenylation must be the 4-OH group of the tyrosine backbone. Compounds **1** and **2** were assigned to their respective chemical shifts based on the ^1^H NMR spectra of L-tyrosine and N-acetyl-L-tyrosine, available on PubChem. Informed by these observations, we proceeded to synthesize chemical standards of 4-O-dimethyllalyltyrosine and its N-acetylated derivative (schemes S1-S2 and S3-S4, respectively) and analyzed them via HRMS and both ^1^H and ^13^C NMR (Fig. S3-S9). The results matched the spectra measured for **1** and **2** from the fungal extract, confirming the chemical identity of the DMATS products.

**Figure 2.**
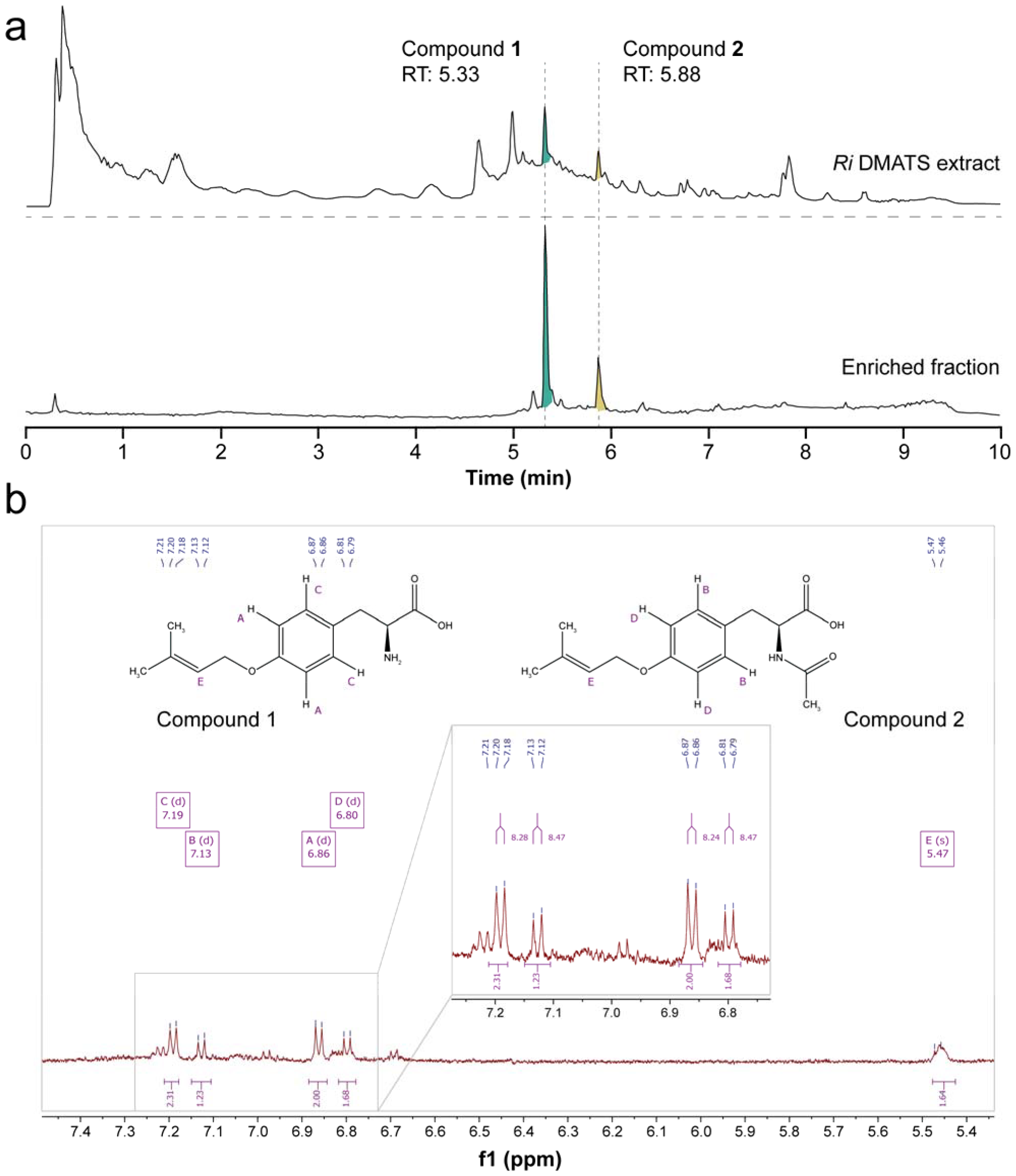
Structural characterization of compounds 1 and 2. **(a)** TICs of total culture extract from Ri DMATS overexpression strain and enriched fraction containing compounds **1** and **2** obtained via semi-preparative HPLC. RTs are different compared to figure 1 because a shorter analysis method was used here; **(b)** Chemical shifts of aromatic protons from ^1^H NMR spectrum (600 Mhz) of enriched fraction. The 4 major doublets indicate that the aromatic rings of **1** and **2** are free of substitutions, consistent with prenylation occurring on the 4-OH group of the substrates. The chemical shift of the prenyl proton is also observed.

### Sequence similarity network analysis suggests that lichen DMATS-type PTs are mainly active on polyketide and phenylpropanoid compounds but not on indoles

To gain further insights into the substrate specificity and activity of lichen DMATS-type PTs, we explored potential relationships between amino acid sequences and substrate specificity across the entire enzyme family. For that, we first collected all sequences of fungal DMATS-type PTs from the InterPro database (IPR012148) [26]. We then manually extracted the sequences of all DMATS-type PT enzymes from 38 lichen genomes available publicly and appended them to cluster (in red) which contains some of the best studied prototypical DMATS enzymes. These are responsible for C4, C5 and C7-prenylation of tryptophan or tryptophan-containing cyclic dipeptides. The indole diterpene (IDT) PTs (light pink), the cyclic dipeptide reverse C2-PTs (fuchsia), and the aromatic polyketide PTs (part of the blue cluster) further exemplify this trend. Furthermore, we carried out a phylogenetic analysis of the biochemically characterized DMATS which revealed that there is a clear distinction between prototype DMATS, PTs active on larger indoles (cyclic peptides and IDTs), and PTs active on non-indole aromatics such as tyrosine, phenylpropnaoids, quinolines and large cyclic polyketides (Fig. S10). This was also observed in a recent publication on a tyrosine O-PT from an edible mushroom [30]. Unsurprisingly, *As* DMATS and *Ri* DMATS cluster closely with the two well characterized 4-O-tyrosine PTs *Lm* SirD and *Cp* TcpD. Interestingly, the majority of DMATS from lichen-forming fungi (Figure 3, black nodes with green outline) reside within this cluster as well. Many of them are found in the vicinity of tyrosine-PTs—including the two enzymes from *Letharia* that we cloned—while the remaining ones are located within the aromatic polyketide PTs subcluster. Two PTs active on quinolinone B, a heterocyclic compound derived from anthranilic acid and O-methyl-tyrosine, are also present in this group. Six lichen PTs appear as singletons, while 7 others form an isolated cluster. The remaining are found in two poorly annotated clusters that contain only one reviewed entry each: the 4-O-dimethylallyltyrosine synthase *An tyr*PT [31] (orange) and a PT active on siccayne (light pink), a phenolic compound likely derived from tyrosine or phenylalanine [32]. These observations strongly suggest that lichen DMATS-type PTs might be mainly active on polyketide compounds and aromatic scaffolds derived from tyrosine and/or phenylalanine, but not on indole-containing scaffolds. In fact, aromatic polyketides (such as depsides, depsidones, chromones, xanthones, and anthraquinones), as well as shikimate-derived phenylpropanoids (terphenylquinolones and pulvinic acid derivatives), are the most abundant natural products isolated from lichen tissues [4,11,33]. In contrast, nitrogen-containing secondary metabolites in lichen are rare and, to the best of our knowledge, indole alkaloids have never been isolated from lichens thus far. This is attributed to the fact that nitrogen is a limiting nutrient for lichen-forming fungi and therefore mostly restricted to primary metabolism [19]. Indeed, our SSN analysis shows that only one out of 67 lichen PTs— from the organism Usnea *florida*—localizes in the prototype DMATS cluster. Interestingly, the corresponding BGC as predicted by fungiSMASH shares a high similarity with the ergotamine cluster from *Claviceps purpurea* (Fig. S11) [34] retrieved from the MIBiG database [35]. Multiple sequence alignment (MSA) also showed that Uf DMATS shares the same catalytic residues of prototype tryptophan DMATS (Fig. S12). Thus, further characterization of the BGC is of great interest as it might lead to the discovery of the first ergot alkaloid biosynthetic pathway in a lichen-forming fungus.

**Figure 3.**
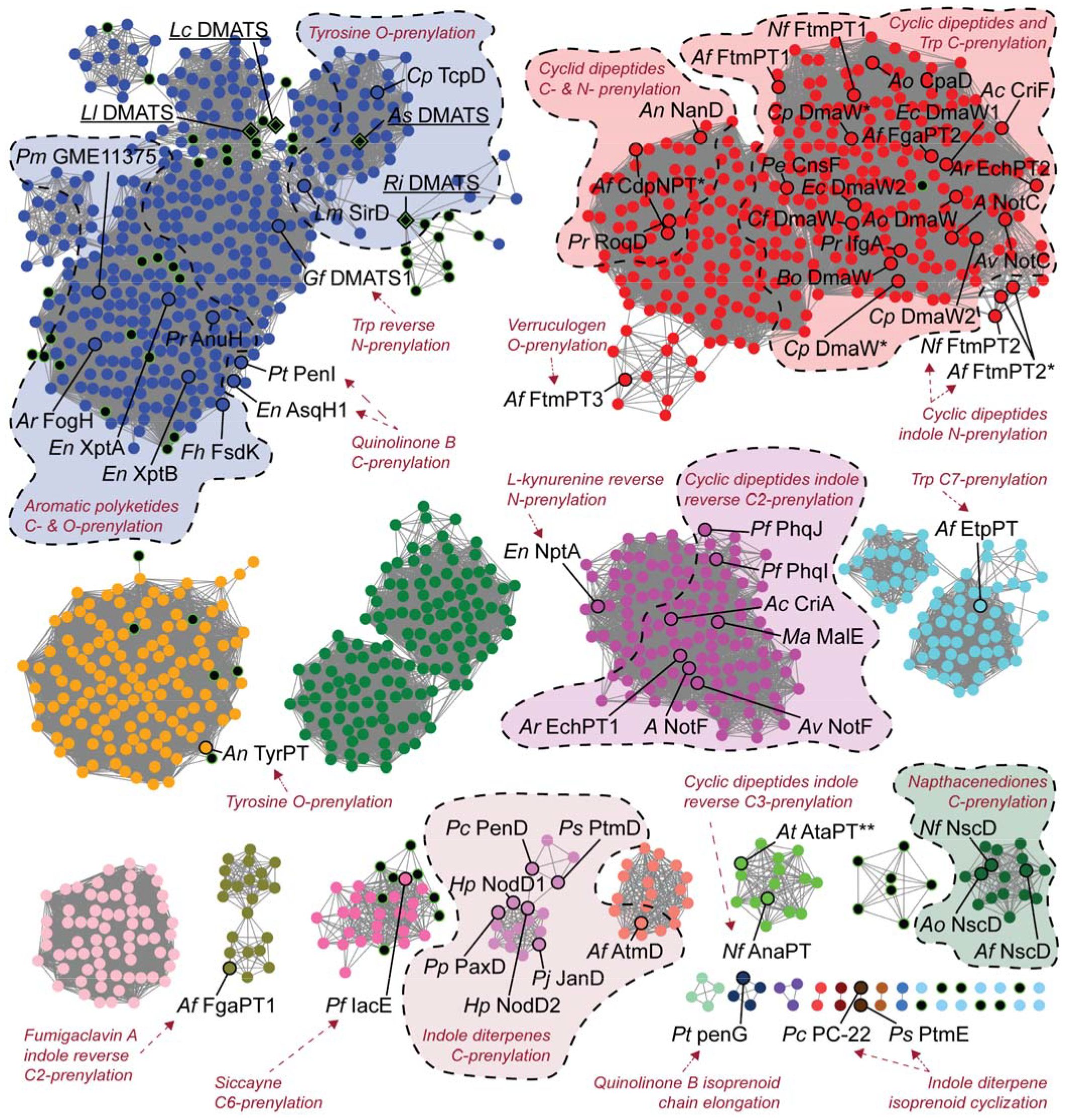
SSN of fungal DMATS-type PTs. The organic layout from yFiles was applied. Edge selection cut-off: sequence ID % > 34.2. The different clusters are colored based on the default color scheme of EFI-EST. Lichen enzymes are colored in black with green borders, re-gardless of their location within the network. The 4 enzymes subbject of this work are highlighted as green diamonds with underlined labels. Enzymes with known substrate specificity and type of reaction catalyzed (indicated in red) are circled in black and labeled (organ-ism + enzyme name). For visualization purposes, dashed lines encircle groups of enzymes that catalyze the same type of reaction, although it should be stated that this notation only refers to characterized enzymes labeled in black. *Enzymes from the same species but different strains. **For simplicity, no annotation on substrate specificity/reaction was included for At AtaPT because of its broad donor (DMAPP, GPP, FPP) and acceptor (peptides, flavonoids, aromatics) substrate scope [35].

### Docking and MSA studies show conserved active site architecture and substrate binding in lichen 4-O-dimethylallyltyrosine synthases

It is well known that DMATS are promiscuous enzymes and some of them can prenylate an incredible range of aromatic substrates *in vitro* (e.g. tyrosine, tryptophan, cyclic peptides, lactones, quinolines, and even plant flavonoids) [36–39]. Though less broad, the tyrosine-PTs *Lm* SirD and *An* TyrPT also show an extended substrate scope which includes tryptophan and derivatives. Furthermore, these enzymes can catalyze prenylation at different positions—and atoms—on the aromatic ring [31,40,41]. Although our experiments were only performed *in vivo*, we found it surprising that both *As* DMATS and *Ri* DMATS overexpression strains only produced 4-O-dimethylallyltyrosine and its N-acetylated derivative. Therefore, we used bioinformatic tools to generate structural models of the enzymes and investigate their active site architecture. First, we used AlphaFold [42] to build models for both As and *Ri* DMATS and superimposed them to the crystal structure of *Af* FgaPT2 [15] to inspect the general structural features (Fig. 4a). The overall ABBA fold and central β-barrel where the reaction chamber is located are very well conserved in the lichen enzymes. The tyrosine shield—a set of four Tyr residues that play a major role in catalysis [17]—also perfectly overlaps with that of FgaPT2. The only differences are observable at the termini of the proteins. At the N-terminus, *Ri* DMATS possesses two extra α-helices (α_1_ and α_2_) compared to FgaPT2, while the N-terminus of *As* DMATS is even shorter than that of FgaPT2, with a short unstructured region in place of helix α_F_. At the C-terminus, the model of *As* DMATS shows a relatively long unstructured region, which is absent in both *Ri* DMATS and FgaPT2. Intrigued by these differences, we also retrieved the structural models of the other 4-O-dimethylallyltyrosine synthases *Lm* SirD, *Cp* TcpD, and *An* TyrPT from the AlphaFold database [43], and observed that all of them possess additional α-helices at their N-termini (Fig. S13). It should be noted that in all cases the structures in this N-terminal region are predicted at very low confidence levels, thus we refrain from formulating any hypothesis on its putative role at this stage.

**Figure 4.**
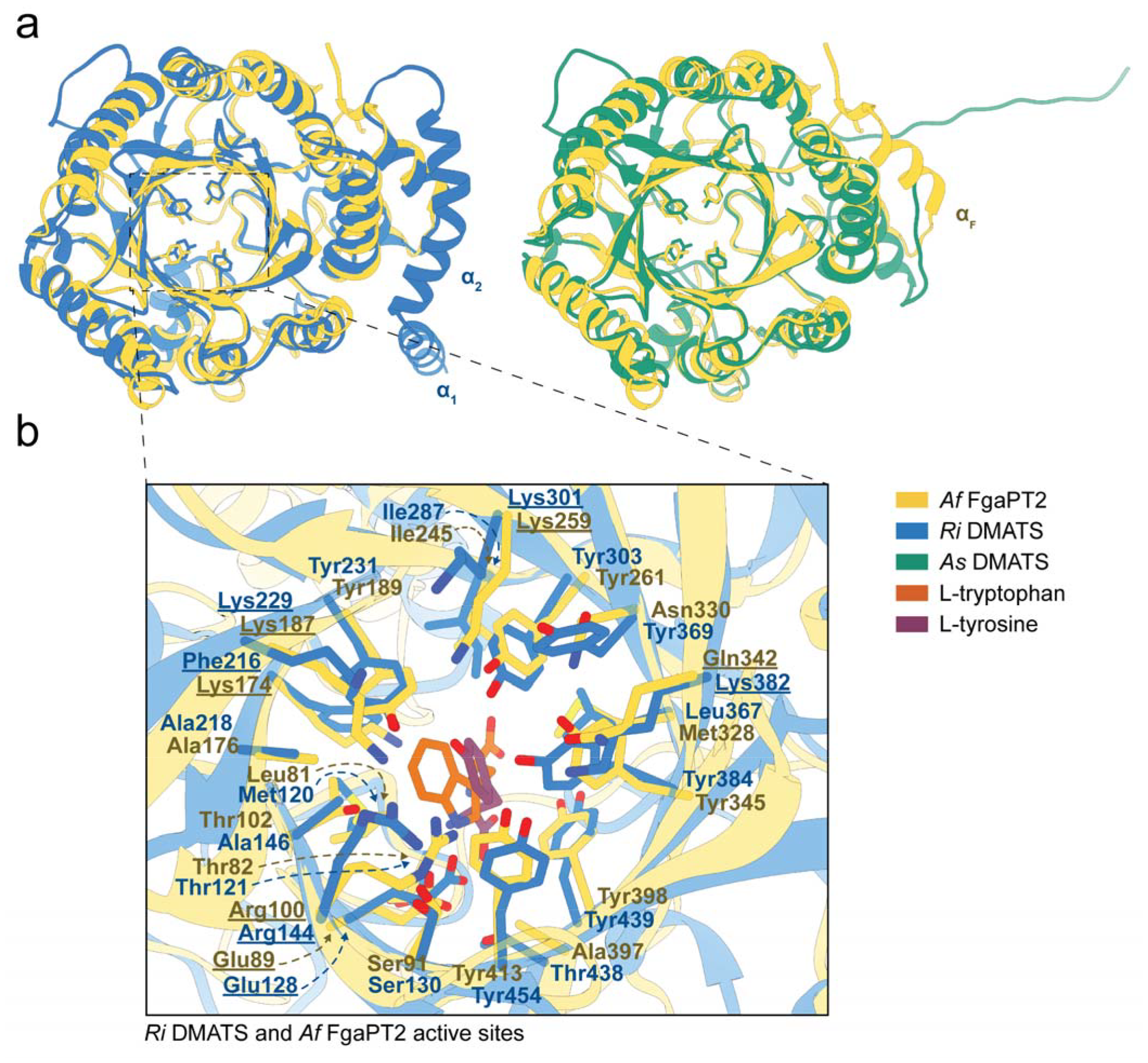
Comparison of *Ri* DMATS and *As* DMATS models with the crystal structure of *Af* FgaPT2 (PDB ID: 3I4X). **(a)** Superimposition of the structures shows that the overall ABBA fold and the central β-barrel where the tyrosine shield and active site are located are well conserved in the lichen DMATS. Two additional α-helices can be observed at the N-terminus of *Ri* DMATS, though these are predicted with low confidence (Fig. S13). In the corresponding region, *As* DMATS is instead missing the N-terminal α-helix present in FgaPT2, and an additional, large, unstructured loop is predicted at its C-terminus (also with low confidence, fig. S13). **(b)** Active site comparison of *Ri* DMATS and *Af* FgaPT2. All amino acids within a 5 Å radius from the substrates are displayed. The binding pockets show a similar architecture and substrate positioning. Key catalytic residues in FgaPT2 and corresponding residues in *Ri* DMATS are underlined. The structures were visualized using UCSF Chimera and aligned with the internal Matchmaker tool.

Next, we decided to have a deeper look into the active site architecture and possible binding mechanism of the lichen DMATS with the substrate L-tyrosine. Given that *As* DMATS and *Ri* DMATS possess the same overall structure and active site residues—and the fact that *Ri* DMATS showed the highest activity in vivo—we focused only on the latter. We first performed rigid molecular docking with the substrates L-tyrosine and DMAPP, followed by a 20 ns MD simulation to achieve reliable energy minimization and “stabilize” the complex. We then superimposed the resulting Tyr-bound structure (DMAPP was hidden for visualization purposes) with the crystal structure of FgaPT2 in complex with its substrate L-tryptophan [15]. The results are shown in Figure 4b.

Unsurprisingly, the overall architecture and the position of the key catalytic residues are well conserved. This is the case for Glu89 (Glu128 in *Ri* DMATS) which interacts with the indole –NH– in FgaPT2 and other prototype DMATS enzymes [17]. It appears that this interaction is retained in *Ri* DMATS, where Glu128 would interact with the –NH_2_ group of L-tyrosine. The DMAPP-binding residues Arg100 (Arg144), Lys187 (Lys229), and Lys259 (Lys301) are also conserved and show the same orientation in the structural models. On the other hand, the key residue Lys174, which acts as catalytic base to deprotonate the intermediate arenium ion in FgaPT2 [15], is replaced by Phe216 in *Ri* DMATS. The aromatic ring of this residue is likely involved in a π–π interaction with the aromatic side chain of the substrate, rather than in catalysis. Indeed, a mutant variant of FgaPT2 where Lys174 was replaced by phenylalanine showed much higher specificity towards L-tyrosine and its activity towards L-tryptophan was almost abolished [44]. Another key difference is Gln343—involved in binding and positioning of the prenyl donor DMAPP in FgaPT2 [15]—which is replaced by Lys382 in *Ri* DMATS. Both these differences are shared by other 4-O-dimethylallyltyrosine synthases (Fig. S12), indicating conserved substrate binding mode and catalytic mechanism. It was recently proposed that *Lm* SirD utilizes the conserved Glu residue (Glu89 in FgaPT2) or a water molecule as catalytic base [17], but our docking model shows that Lys382 might also play a crucial role for 4-O prenylation. In fact, its side chain is positioned in the vicinity of the 4-OH group of L-tyrosine. When we also display DMAPP in the pocket (Fig. S14), it appears clear that the 4-OH group of tyrosine is in a favorable position for the nucleophilic attack onto the dimethylallyl cation. In further support of our hypothesis that Lys382 of *Ri* DMATS has a critical role for 4-O prenylation, the above-mentioned FgaPT2 K174F variant was indeed active on L-tyrosine, but prenylation would occur at the C3 position [44], which in our superimposition almost overlaps with the C4 of L-tryptophan (Fig. 4b). Engineering a double mutant bearing both K174F and Q343K mutations (to the best of our knowledge not yet done) could provide further evidence to validate our hypothesis. Interestingly, when looking at the sequences of fungal aromatic prenyltransferases from other groups than prototype DMATS, we observed that Gln343 is often replaced by lysine, but also arginine residues (and aspartate in one case) (Fig. S12). We also noticed that Lys174 is absent in these sequences, and is replaced by other residues which are conserved across enzymes that cluster together (Fig. S12). As discussed for *Ri* DMATS, all 4-O-dimethylallyltyrosine synthase possess a residue of phenylalanine; aromatic polyketide PTs, cyclic peptide PTs have small hydrophobic residues (Val, Ala, Leu); indole diterpene PTs have methionine residues; and cyclic dipeptide reverse C2-PTs have alanine. Given that these are all nonpolar residues, they are likely not directly involved in catalysis but rather in substrate positioning. The positively charged residues replacing Gln343, instead, might act as catalytic base to deprotonate the reaction intermediates. Interestingly, some indole diterpene PTs and the naphtacenedione PTs are the only ones that have different residues in place of the conserved Glu89 (Fig. S12), which might at least in part have to do with the large size of their substrates.

## CONCLUSIONS

In this work, we report the successful overexpression of two functional DMATS-type PTs, which allowed us to identify them as 4-O-dimethylallyltyrosine synthases. *As* DMATS and *Ri* DMATS are the first enzymes from this family to be described in lichen-forming fungi. Furthermore, we performed SSN, MSA, and phylogenetic analyses which show that all lichen DMATS-type PTs but one are related to known TyrPTs and other aromatic PTs active on polyketides and phenylpropanoids. These are the major metabolites of lichen-forming fungi, which instead do not produce many nitrogen-containing metabolites and are not known to make indole alkaloids. Thus, it is plausible that lichen-forming fungi evolved or maintained only PTs that can diversify non-indole aromatics for enhanced bioactivity. Because many drugs and drug leads bear aromatic polyketides and phenylpropanoids scaffolds [45,46], we believe that further exploring lichen-forming fungi might lead to the discovery of DMATS-type PTs with desirable substrate scopes. These could readily be applied as biocatalysts or in combinatorial pathway engineering, to diversify or functionalize aromatic compounds and improve their biological and pharmacological activities [36,38].

Lastly, we used molecular modeling and docking to investigate the active site architecture of *Ri* DMATS, which revealed that it is shared with known TyrPTs such as *Lm* SirD [40], and likely conserved among all TyrPTs. This suggests that the reaction mechanism is also conserved, though it remains yet to be elucidated experimentally. Generally, it is difficult to accurately predict substrate specificity, prenylation site, and the type of reaction catalyzed by a DMATS-type PT. This is because many factors such as substrates positioning, orientation of catalytic residues, and overall active site architecture, are at play. Nevertheless, we show that by using a combination of bioinformatic tools, it is possible to classify DMATS-type PTs in discrete groups that are active on substrates of the same chemical family. These findings may facilitate prioritization of interesting targets in future efforts aimed at identifying aromatic DMATS-type PTs active on pharmaceutically relevant scaffolds.

## ASSOCIATED CONTENT

**Additional file 1**. Experimental details, supplementary figures and tables for analytical chemistry data and procedures and bioinformatic analyses, synthesis schemes and NMR spectra for reference compounds (PDF).

**Additional file 2**. Full MSA of biochemically characterized fungal DMATS-type PTs (+ target lichen PTs) (FASTA).

**Additional file 3**. SSN file (Cytoscape session) (ZIP).

## AUTHOR INFORMATION

### Author Contributions

All authors have given approval to the final version of the manuscript.

### Notes

The authors declare no competing financial interests.

## ACKNOWLEDGMENTS

The authors are grateful to the staff of the Interfaculty Mass Spectrometry Center of the University of Groningen and University Medical Center Groningen for their services in HRMS and MS/MS analysis. We would also like to thank Prof. Jessica Allen from the Eastern Washington University (Cheney, WA, USA) for the collection of the wild isolates of *Letharia;* Prof. Jun-ichi Maruyama from the University of Tokyo (Tokyo, Japan) for the *A. oryzae* NSAR1 strain; Dr. Colin Lazarus from the University of Bristol (Bristol, UK) for the fungal pTYxxx vectors; Dr. Wonyong Kim from the Korean Lichen Research Institute (Sunchon National University, Suncheon, South Korea) for providing annotations of a number of lichen genomes; Lobke van Goor for performing cultivations of lichen DMATS overexpression strains; and Pieter Tepper for the help with setting up the semipreparative HPLC experiments.

