## Additional file 1 for "Discovery and heterologous expression of functional 4-O-dimethylallyl-L-tyrosine synthases from lichen-forming fungi"

**Contents**

Experimental details 1

Supplementary tables 7

Supplementary figures 10

### EXPERIMENTAL DETAILS

#### Instrumentation and general procedures

$^{1}H$ and $^{13}C$ NMR spectra were recorded on a Bruker Ascend™ 600 Mhz instrument (Billerica, MA, USA) at the NMR facility of the Stratingh Institute for Chemistry, University of Groningen.

HRMS and MS/MS spectra of reference compounds were recorded in positive mode on a Thermo Scientific Q Exactive plus hybrid quadrupole-orbitrap mass spectrometer (Thermo Fisher Scientific, Waltham, MA, USA) at a resolution of 70,000, following direct infusion. Flow rate was set at 100 μl/min and scan range was set at 150-600 m/z. The spectra were inspected and processed using the software MZmine 3 [1].

PCR reactions were carried out using 2x Q5 PCR master mix (New England Biolabs, Ipswich, MA, USA) and 1 μL of template (~10 ng genomic DNA) in a total volume of 25 μL, according to manufacturer's instructions. Primers and other gene-specific parameters are listed in table S3. Standard procedures were used for cloning the assembled constructs in competent *E. coli* DH5α cells. Selection of positive clones was carried using LB agar plates supplemented with ampicillin 100 μg/mL.

#### Construction of expression vectors

The integrative expression vectors pTYargB-eGFPac and pTYadeA-eGFPac were kindly provided by Dr. Colin Lazarus from the University of Bristol, UK. These were digested with FastDigest NotI and PacI and dephosphorylated with FastAP alkaline phosphatase (Thermo Fisher Scientific, Waltham, MA, USA). The ready-to-use vector fragments were subsequently separated from the eGFPac inserts by gel purification using the QIAquick Gel Extraction Kit (Qiagen, Venlo, the Netherlands). The genes encoding *L. lupina* [GenBank acc. no. KAF6225851.1], and *L. columbiana* [GenBank acc. no. KAF6239039.1] DMATS were amplified from the genomic DNA of wild isolates collected in Spokane County, Washington, USA, for previous studies [2]. Preliminary sequencing results showed a frame shift in the putative ORF of *L. columbiana* DMATS. This revealed that the actual, shorter, ORF was almost identical to that of *L. lupina* DMATS (table S4), thus the same primers were used to amplify both genes (table S3). *As* DMATS was amplified from genomic DNA extracted from a culture of *A. strigata* CBS123363 (isolated from limestone in the San Jacinto Mts., San Bernadino County, CA, USA), obtained from the Westerdijk Institute strain collection (Utrecht, The Netherlands). The three genes were cloned by sticky-end ligation in the inducible amyB cassette of the pTYargB vector. The gene encoding *Ri* DMATS was synthesized by Twist Bioscience (South San Francisco, CA, USA) without codon optimization, and further amplified via PCR to add 30bp overlaps for Gibson cloning [3]. The isothermal method (1 h incubation at 50 °C) was used to assemble the insert in the amyB cassette of the pTYadeA vector. Direct-colony PCR was used to pick positive transformants, and the corresponding plasmids extracted using QIAprep Spin Miniprep Kit (Qiagen, Venlo, the Netherlands) were sent to Macrogen Europe (Amsterdam, the Netherlands) for verification by Sanger sequencing.

#### Genetic transformation of *A. oryzae* NSAR1

*A. oryzae* NSAR1 (ΔargB, adeA-, sC-, and niaD-) [4] was kindly provided by Prof. Jun-ichi Maruyama from the University of Tokyo, Japan. The fungus was cultivated on DPY agar plates for routine passages (20 g/L glucose; 10 g/L peptone; 5 g/L yeast extract; 0.5 g/L $MgSO_4 \cdot 7H_2O$; 5 g/L $KH_2PO_4$; microagar 15 g/L in $ddH_2O$; pH 5.5). To achieve profuse sporulation for long-term storage and inoculation of expression cultures, NSAR1 and transformant strains were cultivated on DPY-KCl agar plates (5 g/L glucose; 10 g/L peptone; 5 g/L yeast extract; 0.5 g/L $MgSO_4 \cdot 7H_2O$; 5 g/L $KH_2PO_4$; 45 g/L KCl; 1 mL/L Hutner's trace element solution [5]; microagar 15 g/L in $ddH_2O$; pH 5.5). Spores were harvested by flooding 7-day old colonies with 5 mL of spore harvest solution (8.5 g/L NaCl; 1 mL/L Tween 80 in $ddH_2O$) and manual scraping with sterile loops. Spore solutions were kept at 4 °C for short-term storage or mixed with 45% glycerol 1:2, snap frozen in liquid $N_2$, and kept at -80 °C for long-term storage. Protoplasts of *A. oryzae* were obtained as previously described for other Aspergillus species [6], with some modifications. Briefly, $1x10^8$ conidiospores were inoculated in 25 mL liquid DPY medium and grown overnight (~18h) at 30 °C, 150 rpm. The following day, mycelium was collected by filtration on sterile Miracloth (MilliporeSigma, Burlington, MA, USA), washed with 5-10 mL of fresh DPY medium, and transferred to a clean 50 mL tube. Eight mL of DPY were then added to the mycelium, which was coarsely disrupted with a pipette tip. To this, eight mL of 2x protoplasting solution (1.28 g of VinoTaste Pro (Novozymes, Bagsværd, Denmark) in 10 mL KC buffer

= 82 g/L KCl; 21 g/L citric acid · $H_2O$ in $ddH_2O$; pH 5.8) were added. The total content of the tube was transferred to a clean 125 mL cultivation flask and incubated at 30 °C, 100 rpm. Protoplast formation was monitored every 30 min by examining 10 µL samples under an optical microscope, until a sufficient level was achieved (1.5-2 hours). At this point, 16 mL of 0.6 M KCl were added to the flask, and the mixture was pipetted up and down to release more protoplasts, and filtered again through Miracloth to remove undigested hyphae. The filtrate was centrifuged at 1800g, 10 min, to collect the protoplasts. The supernatant was discarded, and the pellet washed with 2 mL 0.6 KCl and centrifuged at 2400g, 3 min. This procedure was repeated twice. Finally, the pellet was resuspended in 1 mL KTC buffer (45 g/L KCl; 5.5 g/L $CaCl_2$; 10 mM Tris-HCl in $ddH_2O$; pH 7.5) and centrifuged at 2400g, 3 min, one last time. The protoplasts were then resuspended in 600–800 µL KTC buffer, and kept on ice for at least 1 h before proceeding with the following steps. For each transformation, 100 µL of protoplasts were mixed with 1–2 µg (max 10 µL) of the pTY expression vector bearing the lichen DMATS gene (or control eGFP) and 50 µL of PEG-KTC buffer (KTC + 25% w/v PEG4000), and incubated at room temperature for 25 min. After this step, 1 mL of PEG-KTC buffer was added to the mixture, which was incubated on ice for an additional 25 min. Finally, 4 mL of KTC were added, and the transformation mix was centrifuged at 2400g, 3 min. The supernatant was discarded, and the pellet resuspended in 200 µL KTC. These were plated (2 x 100 µL) on 2 TMM agar plates (342.3 g/L sucrose; 2 g/L $NH_4Cl$; 1 g/L $(NH_4)_2SO_4$; 0.5 g/L NaCl; 0.5 g/L KCl; $MgSO_4 \cdot 7H_2O$; 1 g/L $KH_2PO_4$; 1 mL/L Hutner's trace element solution; methionine 1.5 g/L: 1 g/L arginine; 0.1 g/L adenine; microagar 15 g/L in $ddH_2O$; pH 5.5). Arginine or adenine were omitted to select for the vectors bearing the corresponding auxotrophic marker. The protoplasts were regenerated at 30 °C for 5 days, when individual colonies were picked and transferred to new DPY-KCl plates for sporulation and purification of genetically pure clones. For each strain, 4 individual clones were selected and used for the following experiments.

**Cultivation and metabolite extraction**

The DMATS and eGFP (control) overexpression strains were inoculated with sterile cotton sticks from their respective spore suspensions onto MPY agar plates (30 g/L maltose; 10 g/L peptone; 5 g/L yeast extract; 0.5 g/L $MgSO_4 \cdot 7H_2O$; 5 g/L $KH_2PO_4$; microagar 15 g/L in $ddH_2O$; pH 5.5), where maltose was used as inducer for the amyB cassette [7]. The plates were incubated in the dark at 30 °C for 5 days and the control strains checked over a transilluminator on day 3 to confirm eGFP expression and thus, successful induction. For extraction of SMs, the whole agar pads (agar and mycelium) were cut into pieces of roughly 1 $cm^3$ and transferred to 50 mL polypropylene tubes, then extracted once with 25 mL of 9:1 ethyl acetate–methanol (v/v) supplemented with 0.1% formic acid, while being sonicated in a sonication bath for 1 h. Prior to extraction, all samples were spiked with 10 µL of caffeine standard solution (10 mg/mL) to validate the extraction procedure. The organic extracts were collected in clean glass vials and dried under a gentle stream of $N_2$ at room temperature. The dry residues were resuspended in 1 mL of 1:1 MeOH-ultrapure water (v/v) supplemented with 0.1% formic acid by pipetting and vortexing, filtered with 0.45 µm PTFE filters, and stored at -20 °C until further analysis.

**HPLC-MS-DAD analysis of fungal extracts**

HPLC-MS-DAD analysis of the fungal extract was first carried out using a low-resolution Waters Acquity Arc HPLC system coupled to a 2998 PDA detector and a QDa single-quadrupole mass detector (Waters, Milford, MA, USA). A Waters XBridge BEH C18 reversed-phase column was applied for separation (50 mm × 2.1 mm I.D., 3.5 µm, 130 Å particles) which was maintained at 40 °C. The mobile phase consisted of a gradient of solution A (0.1% formic acid in ultrapure water) and solution B (0.1% formic acid in acetonitrile). A split gradient was used: 0–2 min 5% B, 2–10 min linear increase to 50% B, 10-15 min linear increase to 90% B, 15-17 min held at 90% B, 17–17.01 min decrease to 5% B, and 17.01-20 min held at 5% B. The injection volume was 2 µL, and the flow rate was set to 0.5 mL/min. MS analysis was carried out in positive mode, with the following parameters: probe temperature of 600 °C; capillary voltage of 1.0 kV; cone voltage of 15 V; scan range 100–1250 m/z. Data obtained via these experiments was analyzed using the proprietary software MassLynx.

HRMS and MS/MS analyses of fungal extracts were carried out using a Shimadzu Nexera X2 high performance liquid chromatography (HPLC) system with a binary LC20ADXR pump coupled to a Thermo Scientific Q Exactive plus hybrid quadrupole-orbitrap mass spectrometer (Thermo Fisher Scientific, Waltham, MA, USA). A Kinetex EVO C18 reversed-phase column was applied for HPLC separations (100 mm × 2.1 mm I.D., 2.6 µm, 100 Å particles) (Phenomenex, Torrance, CA, USA), which was maintained at 50 °C. The mobile phase consisted of a gradient of solution A (0.1% formic acid in ultrapure water) and solution B (0.1% formic acid in acetonitrile). A linear gradient was used: 0–3 min 5% B, 3–51 min linear increase to 90% B, 51–55 min held at 90% B, 55–55.01 min decrease to 5% B, and 55.01–60 min held at 5% B. The injection volume was 2 µL, and the flow rate was set to 0.25 mL/min. MS and MS/MS analyses were performed with electrospray ionization (ESI) in positive mode at a spray voltage of 3.5 kV, and sheath and auxiliary gas flow set at 60 and 11, respectively. The ion transfer tube temperature was 300 °C. Spectra were acquired in data-dependent mode with a survey scan at m/z 100–1500 at a resolution of 70,000, followed by MS/MS fragmentation of the top 5 precursor ions at a resolution of 17,500. A normalized collision energy of 30 was used for fragmentation, and fragmented precursor ions were dynamically excluded for 10 s. Data obtained by these experiments was analyzed using MZmine 3 [1].

**Large scale cultivation and semipreparative HPLC**

For large scale cultivation and purification of fungal extracts, we inoculated the *Ri* DMATS overexpression strain on eight large 125 mL MPY agar plates (total volume 1 L). The spores were transferred with a sterile cotton stick as before, but this time on 4 equidistant spots on each agar plate. The cultures were then incubated in the dark at 30 °C for 7 days, enclosed in a box to prevent excessive evaporation. For extraction of SMs, the whole agar pads (agar and mycelium) were cut into pieces of roughly 1 $cm^3$ and transferred in a clean borosilicate glass bottle. The total amount of biomass and agar (approximately 1 L) was extracted twice with equal amounts of 9:1 ethyl acetate–methanol (v/v) supplemented with 0.1% formic acid. The extraction mixture was sonicated in a sonication bath for one hour each time. The combined organic extract was collected in a clean borosilicate glass bottle with the help of a separatory funnel. The extract was then concentrated under reduced pressure with a rotary evaporator to yield ~20 mL of concentrated extract. This was transferred to a clean glass vial and dried under a gentle stream of $N_2$ until complete evaporation of the solvents, yielding 989 mg of crude extract. The residue was resuspended in 9.89 mL of 1:1 MeOH-ultrapure water (v/v) supplemented with 0.1% formic acid to achieve a final concentration of ~100 µg/µL, sonicated for 1 h, and filtered with 0.45 µm PTFE filters to remove any undissolved material.

For purification, high-performance liquid chromatography was carried out on a HPLC Shimadzu system, equipped with a dual LC-20AD pump and a SPD-20M20A photodiode array detector (Shimadzu Corporation, Kyoto, Japan). A Macherey-Nagel VarioPrep NucleoDur C18 ec reversed-phase column was applied for separation (250 × 10 mm I.D., 5 µm, 110 Å particles) which was maintained at room temperature (~25 °C). The mobile phase consisted of a gradient of solution A (ultrapure water) and solution B (acetonitrile). Following method development, the following linear gradient was established: 0–15 min 5% B, 15–35 min linear increase to 60% B, 35-40 min held at 60% B, 40–40.01 min decrease to 5% B, and 40.01-60 min held at 5% B. The injection volume was 50 µL, and the flow rate was set to 5 mL/min. Based on comparison between total absorbance chromatograms and total ion chromatograms generated previously, 7 fractions were collected in a scout run, and analyzed on the Waters HPLC-MS-DAD system described above. The mobile phase consisted of a gradient of solution A (0.1% formic acid in ultrapure water) and solution B (0.1% formic acid in acetonitrile). A shorter linear gradient was used: 0–2 min 5% B, 2–5 min linear increase to 90% B, 5–7 min held at 90% B, 7–7.01 min decrease to 5% B, and 7.01–10 min held at 5% B. The injection volume was 2 µL, and the flow rate was set to 0.5 mL/min. Fraction 6 (F6, RT 28.1–28.4 min, Fig. 2a) was determined to be the fraction containing both compounds **1** and **2**. Repeat injections were performed until a total volume of ~40 mL of F6 was collected. The combined fractions were concentrated under reduced pressure to yield an aqueous residue of about 15 mL, which was then snap-frozen in liquid $N_2$ and lyophilized overnight to obtain 2.1 mg dry residue (off-white powder). This was dissolved in 300 µL of $CD_3OD$, transferred to a Norell Select Series™ 3 mm tube (Norell, Morganton, NC, USA) and analyzed via $^1$H NMR (Fig. S2 and Fig. 2b).

**Chemical synthesis of reference compounds**

Diprenyl-(L)-tyrosine: to a suspension of N-Fmoc-L-tyrosine (360 mg, 892 µmol) and $K_2CO_3$ (394 mg, 3.2 equiv.) in $CH_3CN$ (1.8 ml), was added prenyl bromide (247 µl, 2.4 equiv.). The reaction was stirred for 18 hours at RT. Solids were filtered and washed with EtOAc (3x5 ml). The combined filtrates were evaporated to dryness. The crude residue was purified by flash chromatography. Fractions containing spots of Rf = 0.23 in 30% EtOAc in pentanes were combined and evaporated. The reaction afforded 55 mg of the desired product (19%).

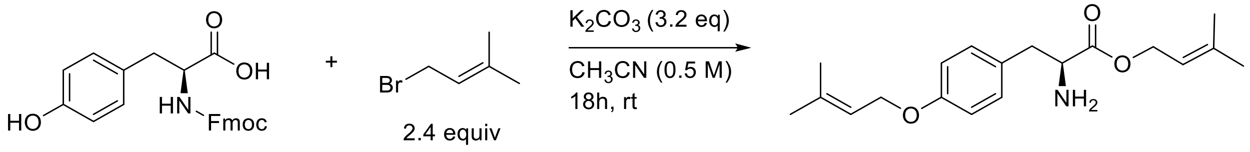


**Scheme S1. Synthesis scheme for diprenyl-(L)-tyrosine.**

4-O-prenyl-(L)-tyrosine: diprenyl-(L)-tyrosine (55 mg, 173 µmol) was suspended in water (0.8 ml), and LiOH.$H_2O$ (4.1 mg, 1.0 equiv.) was added. THF (0.8 ml) was added, and the resulting milky mixture was stirred overnight. TLC showed complete conversion. The volatiles were evaporated, and the residue was azeotrope with toluene (5x5 ml) until solid. The solid was triturated with MTBE (2x5 ml) and pentane (5 ml). Reaction afforded 21 mg (48%) of white solid.

$^1$H NMR (400 MHz, $CD_3OD$) δ 7.21 – 7.12 (m, 2H), 6.87 – 6.79 (m, 2H), 5.44 (tdd, *J* = 6.8, 2.9, 1.5 Hz, 1H), 4.49 (d, *J* = 6.6 Hz, 2H), 3.45 (dd, *J* = 8.2, 4.6 Hz, 1H), 3.06 (dd, *J* = 13.7, 4.6 Hz, 1H), 2.74 (dd, *J* = 13.7, 8.1 Hz, 1H), 1.77 (s, 3H), 1.74 (s, 3H).
$^{13}$C NMR (101 MHz, $CD_3OD$) δ 159.03, 138.34, 131.63, 131.40, 121.42, 115.79, 65.80, 58.85, 41.39, 25.84, 18.16.
HRMS-ESI+ (*m/z*): $[M+H]^+$ calculated for 250.1443; found, 250.1437.

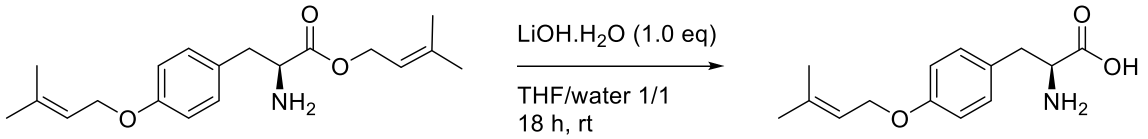


**Scheme S2. Synthesis scheme for 4-O-prenyl-(L)-tyrosine**

4-O-prenyl-N-acetyl-(L)-tyrosine ethyl ester: N-acetyl-(L)-tyrosine ethyl ester monohydrate (125.6 mg, 500 µmol) was suspended in $CH_3CN$ (1.0 ml). $K_2CO_3$ (172.7 mg, 2.5 equiv.) followed by prenyl bromide (69.2 µl, 1.2 equiv.) were added and the resulting suspension was stirred for two days at RT. The reaction mixture was diluted with EtOAc (20 ml), washed with water (2x10 ml) and brine (10 ml), filtered over a phase separator, and evaporated to dryness. The reaction afforded 158.0 mg of a viscous liquid that was used without further manipulation.

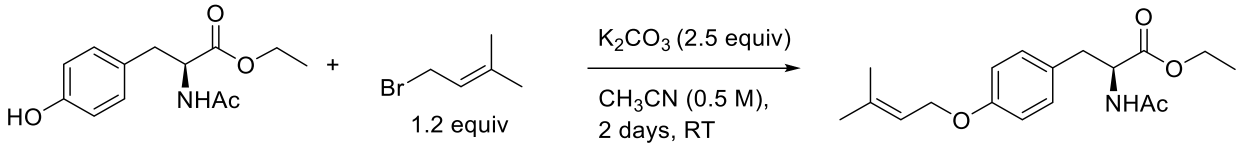


**Scheme S3. Synthesis scheme for 4-O-prenyl-N-acetyl-(L)-tyrosine ethyl ester.**

4-O-prenyl-N-acetyl-(L)-tyrosine: 4-O-prenyl-N-acetyl-(L)-tyrosine ethyl ester (159.7 mg, 500 µmol) was suspended in water (2.5 ml), and LiOH.$H_2O$ (21.0 mg, 1.0 equiv.) was added. The reaction was stirred overnight. After TLC showed the complete conversion, the volume was reduced to ca. 20% by rotavap. The residue was diluted with water (5 ml), transferred into a separatory funnel and extracted with MTBE (3x10 ml). The extracts were discarded. The aqueous phase was acidified to pH = 2 by careful addition of HCl (ca. 1 M, ca. 800 µl). The aqueous phase was extracted with EtOAc (3x10 ml). Combined extracts were washed with brine, filtered over a phase separator and evaporated to dryness. Reaction afforded 140 mg (76%) of white waxy solid. $^1$H NMR suggests an EtOAc solvate (ca. 1/1 desired product/EtOAc).

[1]H NMR (400 MHz, $CD_3OD$) δ 7.12 (d, *J* = 8.6 Hz, 2H), 6.82 (d, *J* = 8.6 Hz, 2H), 5.44 (ddt, *J* = 6.6, 5.2, 1.4 Hz, 1H), 4.60 (dd, *J* = 8.8, 5.1 Hz, 1H), 4.49 (d, *J* = 6.4 Hz, 2H), 4.10 (q, *J* = 7.1 Hz, 2H, EtOAc), 3.12 (dd, *J* = 14.0, 5.1 Hz, 1H), 2.87 (dd, *J* = 14.0, 8.9 Hz, 1H), 2.01 (s, 3H, EtOAc), 1.99 (s, 1H), 1.91 (s, 3H), 1.78 (s, 3H), 1.73 (d, *J* = 1.3 Hz, 3H), 1.24 (t, *J* = 7.1 Hz, 3H, EtOAc).
[13]C NMR (101 MHz, $cd_3od$) δ 174.90, 173.10, 172.98 (EtOAc), 159.21, 138.43, 131.18, 130.37, 121.33, 115.69, 65.81, 61.53 (EtOAc), 55.38, 37.66, 25.84, 22.31, 20.85 (EtOAc), 18.16, 14.46 (EtOAc).
HRMS-ESI+ (*m/z*): $[M+H]^+$ calculated for 292.1549; found, 292.1541.

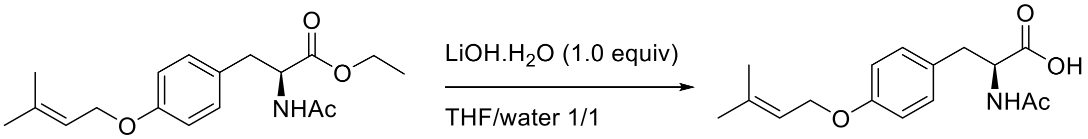


**Scheme S4. Synthesis scheme for 4-O-prenyl-N-acetyl-(L)-tyrosine.**

#### Sourcing putative lichen DMATS sequences

Annotated genomes of 21 lichen-forming fungi were kindly provided by Dr. Wonyong Kim from the Korean Lichen Research Institute, Sunchon National University, Suncheon, South Korea [8]. The remaining 17 were retrieved from the NCBI and JGI Mycocosm databases [9,10], accessed in Septmber, 2021. The annotated genomes were used as input for the biosynthetic gene cluster prediction by antiSMASH v7.0.0 [11] (parameters: --taxon fungi --cassis --clusterhmmer --genefinding-tool none). Of the 2038 clusters identified, 60 were predicted to contain a DMATS-type prenyltransferase. From there, 61 putative lichen DMATS were retrieved.

#### Sequence similarity network analysis

All protein sequences that are part of the fungal aromatic prenyltransferase DMATS-type family (IPR012148) were retrieved from the InterPro database [12] and combined with the DMATS sequences mined manually from lichen genomes, yielding a total of 1446 unique sequences. In total, 67 sequences originate from lichen organisms: 6 were already present in the InterPro dataset and 61 were added based on our manual search, described above. The resulting multi-FASTA file was used as input for generating a sequence similarity network with the Enzyme Similarity Tool of the Enzyme Function Initiative (EFI-EST) [13]; edge selection cut-off: alignment score > 82 (corresponding to sequence ID > 34.23%). The network was colored using the Color SSN utility and visualized in Cytoscape v3.8.2 [14] with the yFiles organic layout.

#### Multiple sequence alignment analysis

The MSA analysis of biochemically characterized fungal DMATS-type PTs and target lichen PTs (and *Uf* DMATS) was performed using MEGA 11 [15] using the MUSCLE algorithm [16] with default settings. The resulting alignment was exported in FASTA format and used as input for the phylogenetic analysis. The software IQ-TREE v2.2.2.6 [17] was used to build a consensus phylogenetic tree with the bootstrap method using the following parameters: --seqtype AA -m TEST -b 1000 (number of replicates = 1000). The final tree was visualized and refined with the online tool Interactive Tree Of Life (iTOL) v6 [18].

#### Modelling and molecular docking

AlphaFold2 Colab was utilized to predict the structure of *Ri* DMATS and *As* DMATS based on their amino acid sequences [19,20], followed by an additional relaxation step employing Amber [21] to ensure precise positioning of the residue side chains. Docking of

DMAPP and L-tyrosine into the active site of the generated *Ri* DMATS model was carried out using AutoDock Vina [22,23]. Following this, a 20 ns molecular dynamics (MD) simulation was performed using YASARA v23.5.19 (YASARA Biosciences). For that, the YASARA dynamics software package incorporating the YAMBER3 force field was used [24]. Initially, the enzyme–substrate complexes were cleaned, minimized, and placed within a rectangular simulation cell, with the distances between the protein and the periodic boundaries of the cell maintained at a minimum of 7.5 Å. The Berendsen thermostat algorithm [25] and manometer algorithm were used to maintain consistent temperature and pressure values during MD simulations. The physiological conditions of the simulation cells were set to mimic 298 K, pH 7.4, and 0.9% NaCl. Temperature was gradually increased from 5 to 298 K over 30 ps, followed by a 4970 ps equilibration period before the production phase simulation of 20,000 ps. Snapshots were captured every 25 ps. Finally, snapshots of the DMAPP and L-Tyr–bound *Ri* DMATS model were captured at the end of the simulation to analyze the active site architecture and binding of the substrates.

**Visualization and other bioinformatic tools**

For visualization of molecular models and docking, UCSF Chimera v1.17.3 was used [26]. Custom settings were used to display ribbon and atoms in cartoon style, and flat lighting was applied. Variations of the Wong and Tol colorblind-accessible palettes were used. Clinker [27] was used, with standard settings, to display and annotate the BGC containing the putative prototype DMATS from *Usnea florida* shown in Fig. S11. Jalview [28] was used to visualize and generate the MSA alignment shown (partially) in Fig. S12. CLUSTAL coloring was applied to the amino acid sequence.

**Table S1. Genomic location of target DMATS from lichen-forming fungi.**

| Gene | Organism & genome assembly | Scaffold/contig | region ID (fungiSMASH) | Location BGC and gene (nt) |
|---|---|---|---|---|
| *As* DMATS | *Acarospora strigata* CBS132363 [29] – JGI Mycocosm database [9] | scaffold 8982 | AS-522.1 | 43,412 – 87,968 (58,206 – 61,127) |
| *Ri* DMATS | *Ramalina intermedia* YAF0013 – unpublished – GenBank acc. no. GCA_003073195.1 | scaffold 38 (acc. no. PEKF01000038.1) | RI-146.1 | 107,028 – 128,413 (117,028 – 118,413) |
| *Ll* DMATS (GenBank acc. no. KAF6225851.1 | *Letharia lupina* WasteWater1 isolate [30] – GenBank acc. no. GCA_014066315.1 | contig 7 (acc. no. JACCJB010000007.1) | LL-28.2 | 2,058,775 – 2,089,139 (2,077,949 – 2,079,139) |
| *Lc* DMATS (GenBank acc. no. KAF6239039.1) | *Letharia columbiana* WasteWater2 isolate [30] – GenBank acc. no. GCA_014066305.1 | contig 7 (acc. no. JACCJC010000007.1) | LC-7.2 | 767,644 – 791,438 (777,644 – 781,438) |

**Table S2. Comparison of HRMS and MS$^2$ data of compound 1 and 2 (from total fungal extract) with synthetic standards. MS$^2$ spectra were recorded in DDA mode (top 5 peaks per MS$^1$ scan).**

| Compound | Molecular formula | $[M+H]^+$ precursor (error, ppm) | Top MS/MS fragment ions[a] | |
|---|---|---|---|---|
| | | | $[M+H]^+$ | Intensity, %[b] |
| 1 | $C_{14}H_{19}NO_3$ | 250.1438 (-2.00) | 165.0544 | 100 |
| | | | 136.0765 | 52 |
| | | | 69.0698 | 26 |
| | | | 123.0447 | 23 |
| | | | 147.0437 | 19 |
| | | | 119.0491 | 11 |
| 2 | $C_{16}H_{22}NO_4$ | 292.1547 (-0.68) | 204.1384 | 100 |
| | | | 136.0765 | 100 |
| | | | 246.1495 | 54 |
| | | | 178.0868 | 53 |
| | | | 233.1171 | 47 |
| | | | 182.0816 | 39 |
| | | | 165.0543 | 29 |
| | | | 69.0704 | 20 |
| | | | 250.1456 | 18 |
| PFO163 | $C_{14}H_{19}NO_3$ | 250.1437 (-2.40) | 165.0541 | 100 |
| | | | 136.0753 | 46 |
| | | | 69.0701 | 25 |
| | | | 123.0438 | 21 |
| | | | 147.0438 | 17 |
| | | | 119.0495 | 11 |
| PFO173 | $C_{16}H_{22}NO_4$ | 292.1541 (-2.74) | 136.0752 | 100 |
| | | | 204.1397 | 84 |
| | | | 178.0862 | 68 |
| | | | 182.0809 | 49 |
| | | | 246.1506 | 41 |
| | | | 233.1184 | 40 |
| | | | 165.0540 | 32 |
| | | | 69.0699 | 28 |
| | | | 250.1422 | 15 |
| | | | 205.1420 | 10 |
| | | | 137.0789 | 10 |

a. Only fragment ions with % intensity > 10 are shown
b. Peak height relative to main fragment

**Table S3. Primers used to clone target lichen DMATS.**

| Target | FW primer[a] (5' → 3') | RV primer[a] (5' → 3') | $T_{ann}$ | Extension time |
|---|---|---|---|---|
| *Ll* DMATS<br>*Lc* DMATS | *ctag*GCGGCCGCatgataggtcgc<br>cctctagctctg | *tagc*TTAATTAActatgctttccatct<br>agcaacatgatagatttc | 67 °C | 30 sec |
| *As* DMATS | *atcg*GCGGCCGCatgagttgcgga<br>ggtaacatggac | *atgc*TTAATTAAttaagactgtgac<br>gctttcatgtgc | 68 °C | 30 sec |
| *Ri* DMATS | **ctcccttctctgaacaataaaccccaca**<br>**gc**gcggccgcatggctggtgaggccagt<br>caaatg | **cagtaccatcatatactctccacccttaa**<br>**t**ttatgcataggcctttggagcaatataaga<br>tgtcaat | 72 °C | 35 sec |

a. 4-bp cleavage overhangs are italicized; recognition sites for restriction enzymes NotI (FW) and PacI (RV) are capitalized; 30-bp overlap regions for Gibson cloning of *Ri* DMATS are underlined and in bold.

NB: *L. lupina* and *L. columbiana* are closely related, and the corresponding target DMATS are homologs and display 98.99% amino acid sequence identity.

**Table S4. Amino acid sequences of target lichen DMATS.**

| Enzyme | Length | Sequence |
|---|---|---|
| *Ll* DMATS | 396 | MIGRPLALLLHEAGYDIHNQYGSLLFFRHCIAGRLGARPTSTGSPQVWKSFMTDDFSPVEYS<br>WCWDTPKGPPRIRFSVDAIGPDAGTQSDPFNQEMTTDLVRHVESVASNVDWKLFNHFRNA<br>FCEQGLEKRVSEGCDDLEKSHTSSIFMAFELHKSEVAVKAYFVPVKAEQTGRSRLSVLSDSIVS<br>LEKSDLRVGAYDQMLAFMTSDAEGSHLEIVGIAVDCVLPKDSRLKLYVRSPSTSFDSVCAIMT<br>LGGKLNTFPQATLKDFRKLWQLTLGLGEDFAPGANLQAKSHETAGVLYNFDIKAGNLLPEP<br>KVYIPVRHYARNDLAAAEGLASYLKSKGQDRFVESYMRALEGMCTHRFLGSQCGLQTYISC<br>AVQNAQLVLTSYLSPEIYHVARWKA |
| *Lc* DMATS | 396 | MIGRPLALLLHEAGYDIHNQYGSLLFFRHCIAGRLGARPTSTGSPQVWRSFMTDDFSPVEYS<br>WCWDTPKGPPRIRFSVDAIGPDAGTQSDPFNQEMTTDLVRHVESVASNVDWKLFNHFRNA<br>FCEQGLEKRVSEGCDDLEKSHTSSIFMAFELHKSEVAVKAYFVPVKAEQTGRSRLSVLSDSIVS<br>LEKSDLRVGAYDQMLAFMTSDAEGSHLEIVGIAVDCVLPKDSRLKLYVRSPSTSFDSVCAIMT<br>LGGKLNTFPQATWKDFRKLWQLTLGLGEDFAPGANLQAKSHETAGVLYNFDIKAGNLLPEP<br>KVYIPVKHYARNDLAAAKGLASYLKSKGQDRFVESYMRALEGMCTHRFLGSQCGLQTYISC<br>AVQNAQLVLTSYLSPEIYHVARWKA |
| *As* DMATS | 435 | MSCGGNMDGLDAGEALNSLSWQNSAGVMLSEMMEMAGYHLQSQRSHLDFFARHVAPAL<br>GSHPEIDRKPPPWRSFMTDDGSPIELSWSWSVQEPAPIVRYSIEPIGDRAGLCPDYFNTHTSN<br>ELVHIIQRSYQGVDLTGYAHFFKELVVCGEGTFVPKITKEDGSNSQIFLAFDLLDEKIMLKVY<br>FLPALRARETGQCKLSMVEKAISTLSPHGQSLSGAFSLVCEYIRSLKVGNRPEIEIIAVDCVNPT<br>LSRVKVYLRSRETSFASVVSMMTLGGRLKELSKEGFATLEELWRLVLSLDPTISTSEPLHLNNR<br>RTAGILYYFELQPSRSYPKPKVYIPVKHYGKSDLKVANGLSSYLKEKGKRLNGMDYRDALQRL<br>CKHRPLDQGSGLHTYVACAIEDISLAVTAYINPEIYHRPRPANRAPAHRETSHMKASQS |
| *Ri* DMATS | 461 | MAGEASQMLRSKRDSMSWQHQYERLASARSELLKRVQLGETVTSTPKIWELITSLQLSIDED<br>VRFWWTVLGTPLAILFQKAGYSIESQYQHLLFFYFLVAPELGARSDGQGLPTTWKSFMTDH<br>FTPIEMSWEWGSNSDGGPTIRYAFEPISAHAGTTLNPLNEGASTRVMHRYSQMIPGCDMTL<br>FHHFAQDLLCYDPSPVSTQGKINSQGHASRCFLAIEFNKSEVMVKAYFFPTFKAIRTNQCPW<br>TMISESILNMPGYSSLMLQSLSSFETFLRCSPEGLNLVPEILAIDCGPPAESRMKIYMRSRSTTF<br>ESVRRVMTLDGALRESGLEKGLYELYVLWTLVFWHGRQVAPEASLQSVEHRTAGILYYFNLS<br>QGGQPPSVKVYLPVRHYGYSDCDVAQGVITYLRSRGRSCSTTEYIEALTAIARPKSLGSQRGL<br>QTYLGCSIVGEKLKLTSYIAPKAYA |

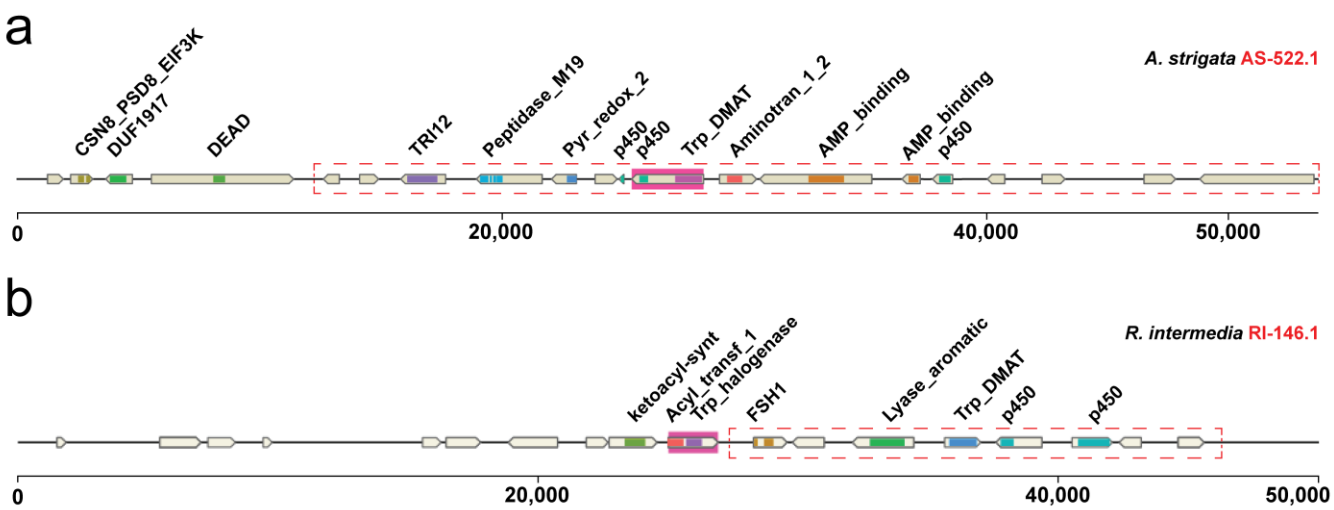


**Figure S1. Genome neighborhoods of target DMATS from *A. strigata* and *R. intermedia*.** The genomic regions were visualized with the gggenomes R package [31]. The specific PFAM domain hits are highlighted by colors and corresponding labels. Corresponding BGCs as predicted by fungiSMASH [11] are highlighted by red dashed boxes. **(a)** Neighborhood (25kb up- and downstream) of putative di-domain DMATS-P450 from *A. strigata*, highlighted in pink. Sequencing results later revealed that this was, in fact, not a fusion protein. **(b)** Neighborhood (25kb up- and downstream) of putative di-domain acyltransferase-halogenase from *R. intermedia*, highlighted in pink. The DMATS gene subject of this study co-localizes downstream of the di-domain protein and is annotated as Trp_DMAT.

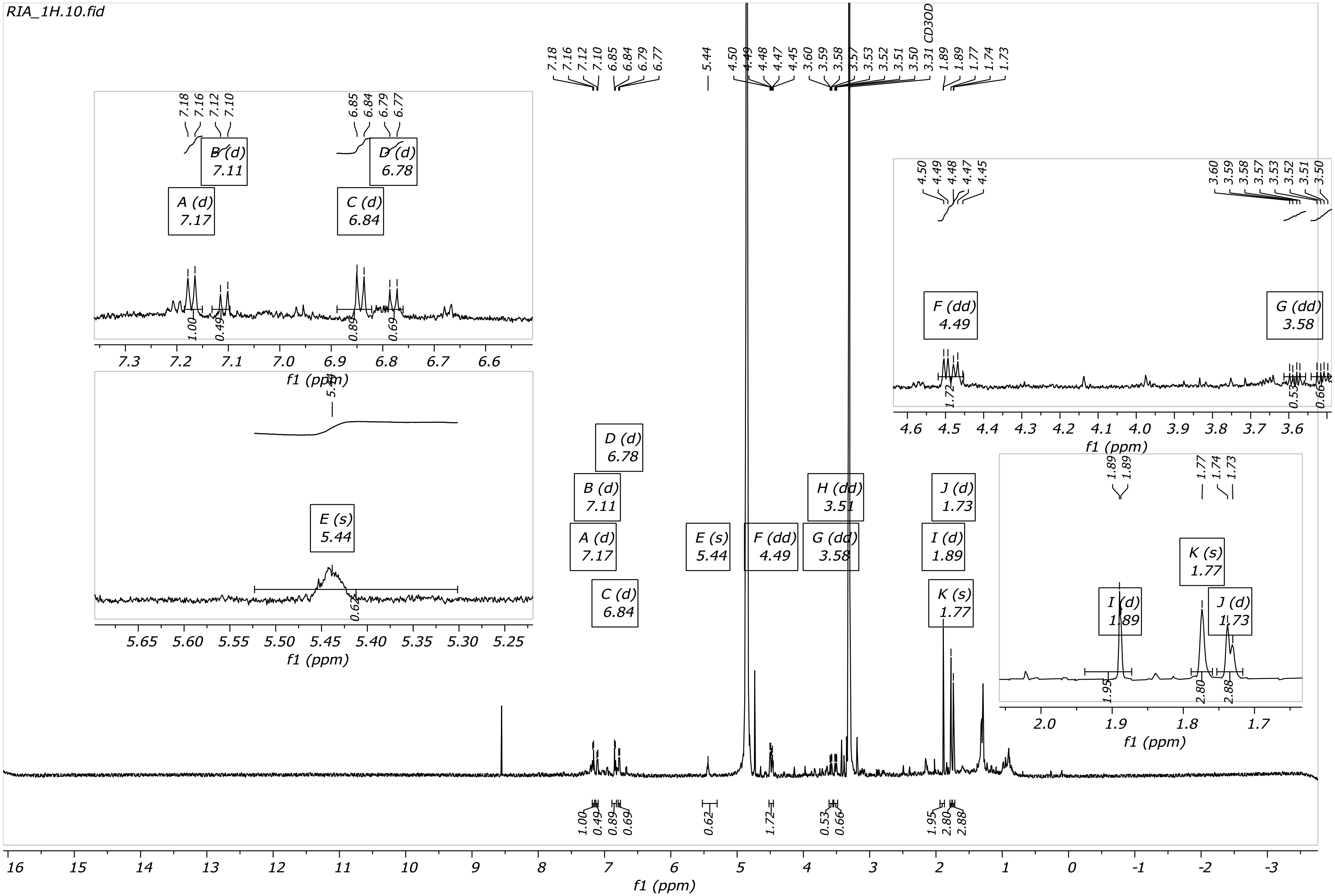


**Figure S2.** ¹H NMR spectrum (600 Mhz) of enriched fraction from *Ri* DMATS extract, containing compound 1 and 2, in $CD_3OD$.

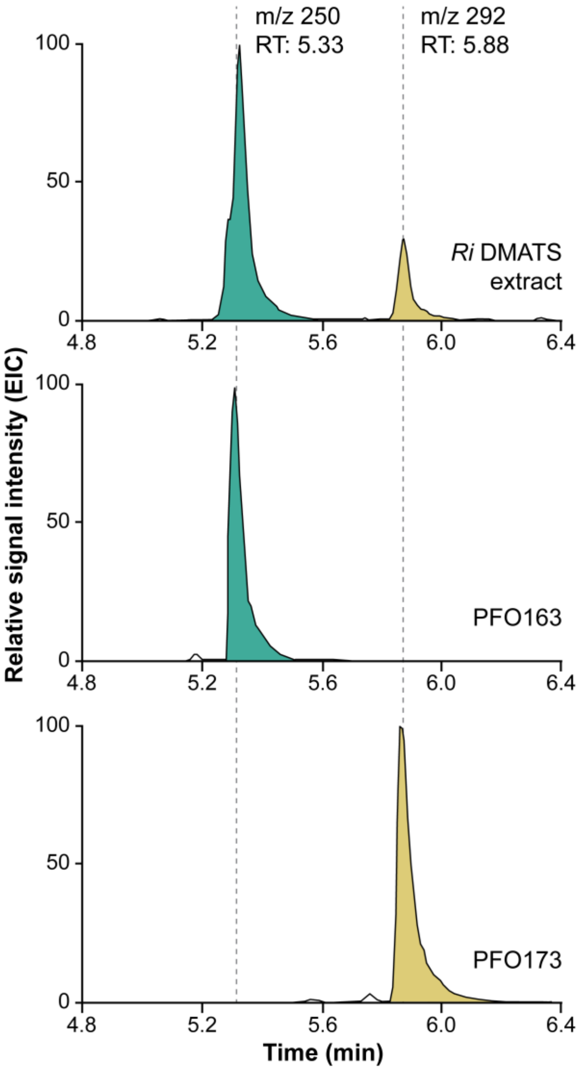


**Figure S3. Comparison of extracted ion chromatograms of the enriched fraction from *Ri* DMATS extract and reference standards PFO163 and PFO173.** Filters were set at m/z 250 (± 0.2 Da) and 292 (± 0.2 Da) for compound **1** and **2** and their references, respectively, in positive ion mode (ESI+).

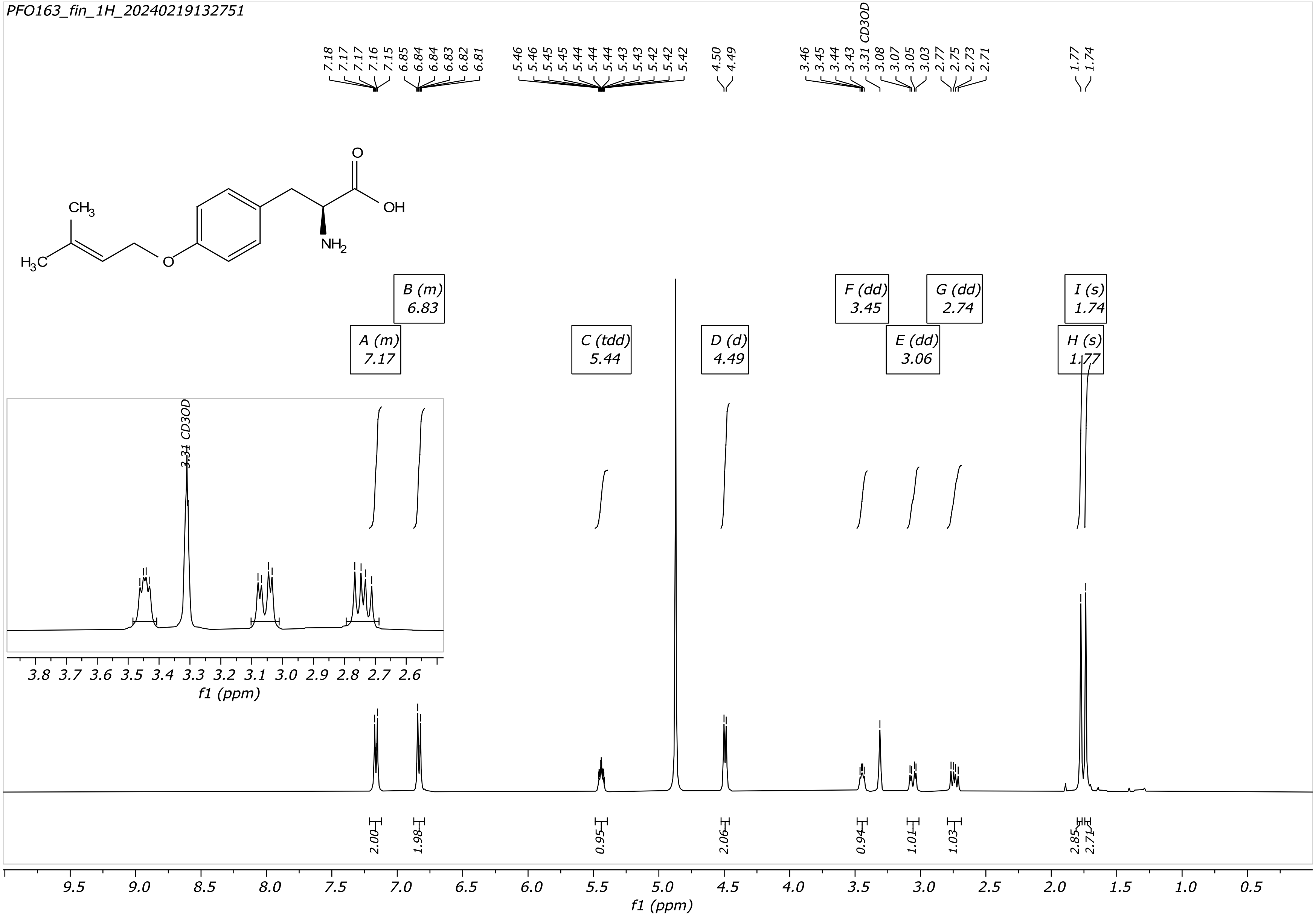


**Figure S4.** [1]H NMR spectrum (600 Mhz) of compound PFO163: 4-O-dimethylallyl-L-tyrosine, in $CD_3OD$.

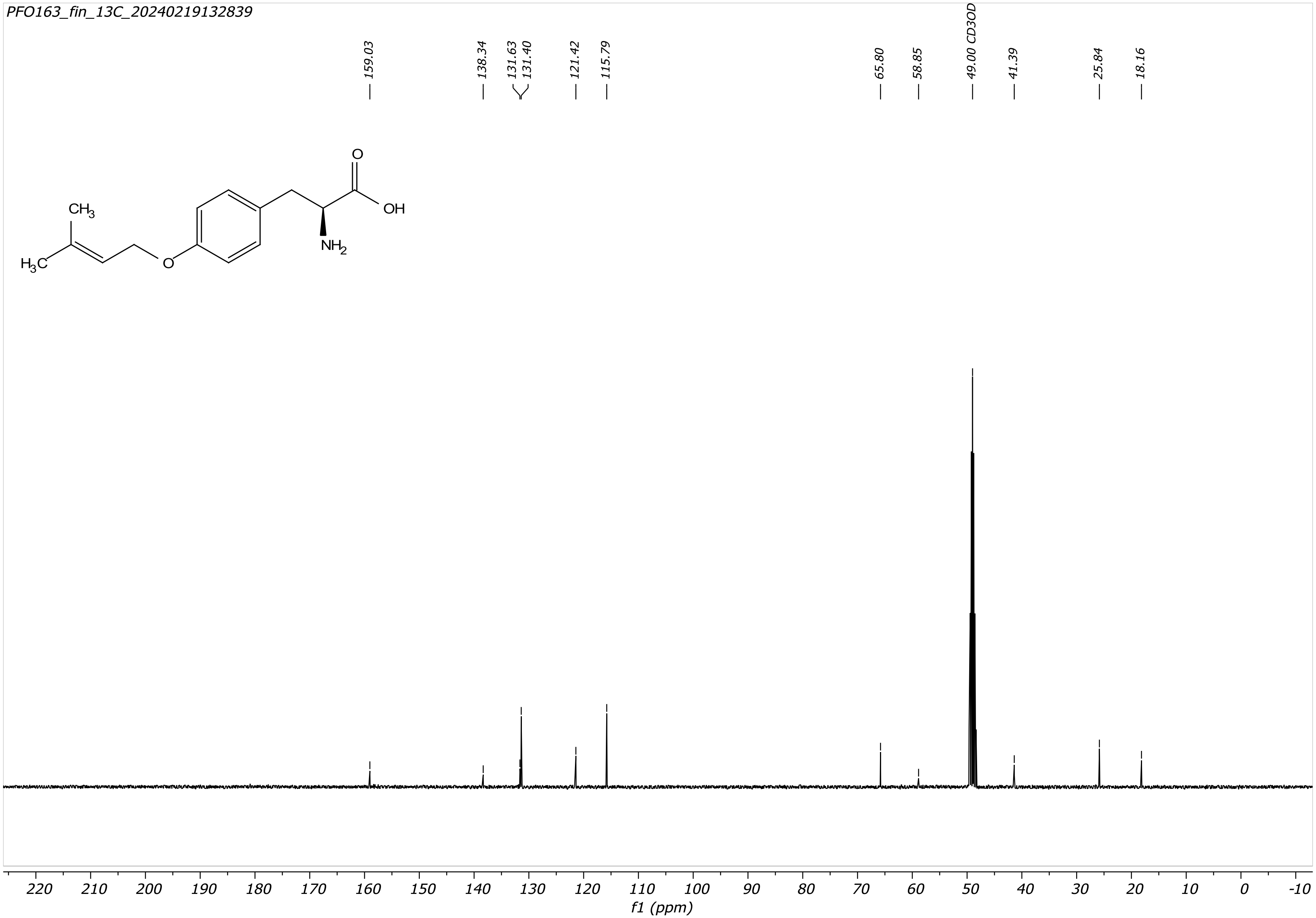


**Figure S5.** **$^{13}C$ NMR spectrum (150 Mhz) of compound PFO163: 4-O-dimethylallyl-L-tyrosine, in $CD_3OD$.**

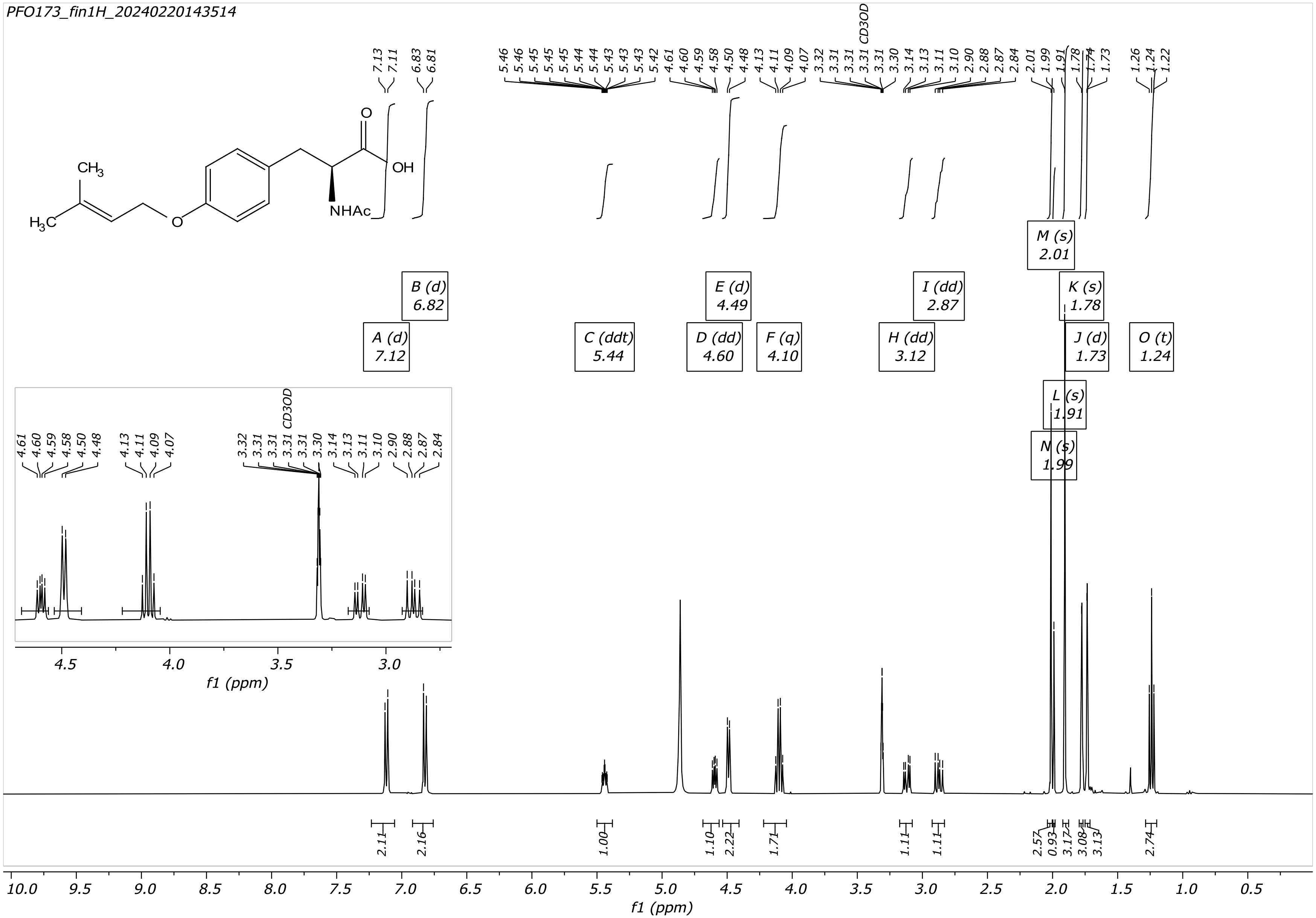


**Figure S6.** **¹H NMR spectrum (600 Mhz) of compound PFO173: 4-O-dimethylallyl-N-acetyl-L-tyrosine, in $CD_3OD$.**

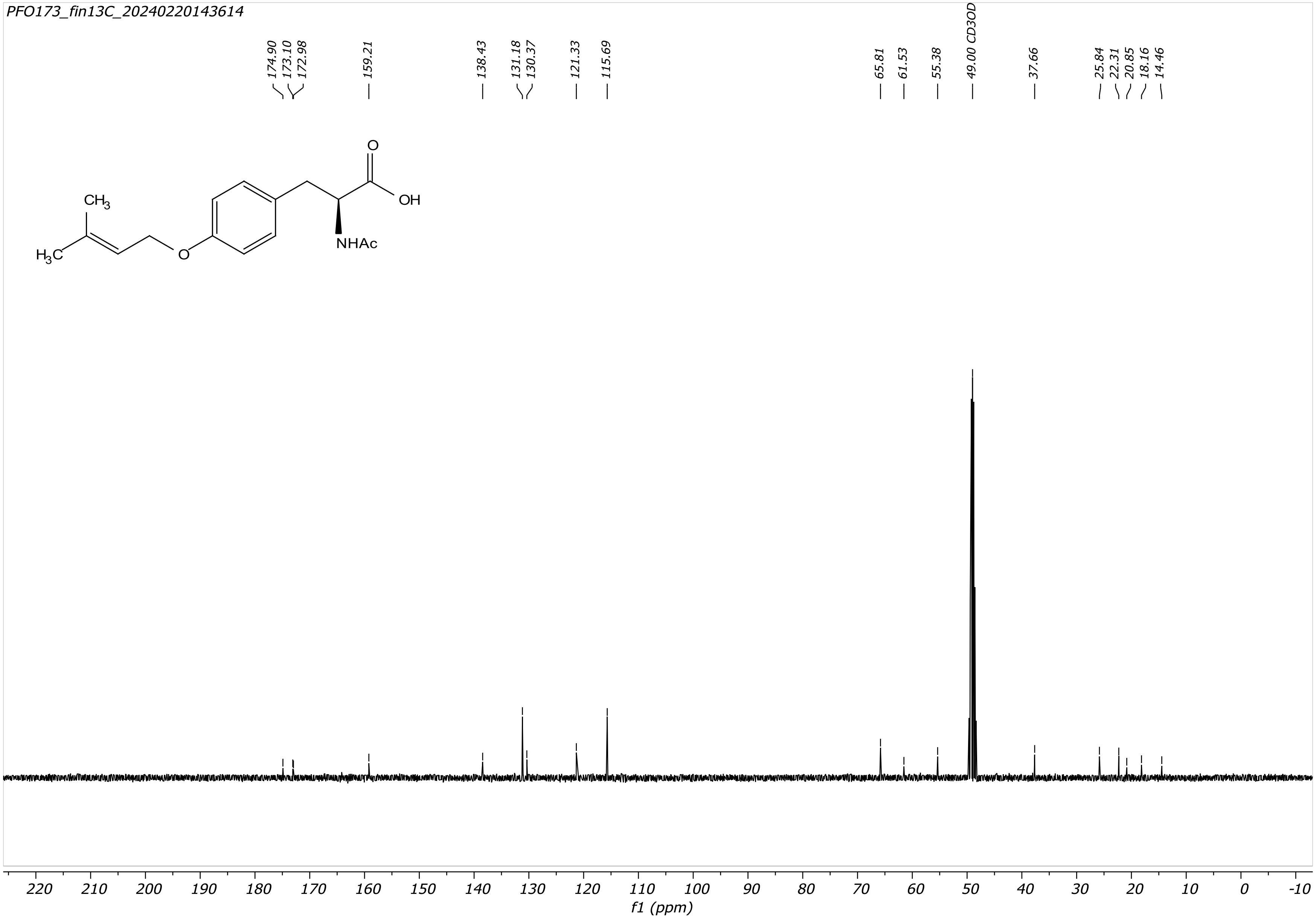


**Figure S7.** $^{13}$C NMR spectrum (150 Mhz) of compound PFO173: 4-O-dimethylallyl-N-acetyl-L-tyrosine, in $CD_3OD$.

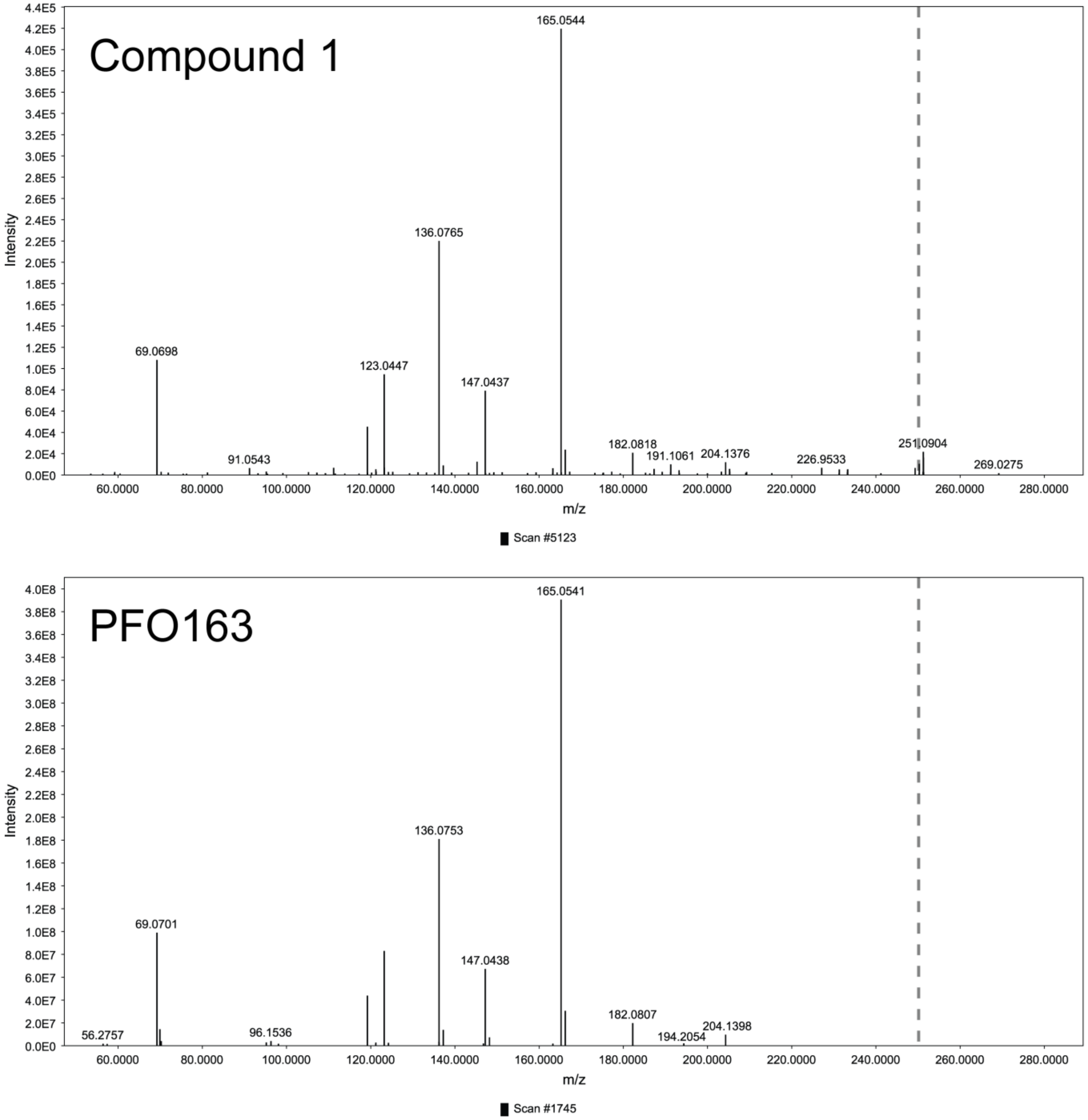


**Figure S8. Comparison of $MS^2$ spectra of compound 1 from the extract of the *Ri* DMATS overexpression strain and its chemically-synthesized reference compound, PFO163.** Precursor m/z value is indicated by a dashed grey line.

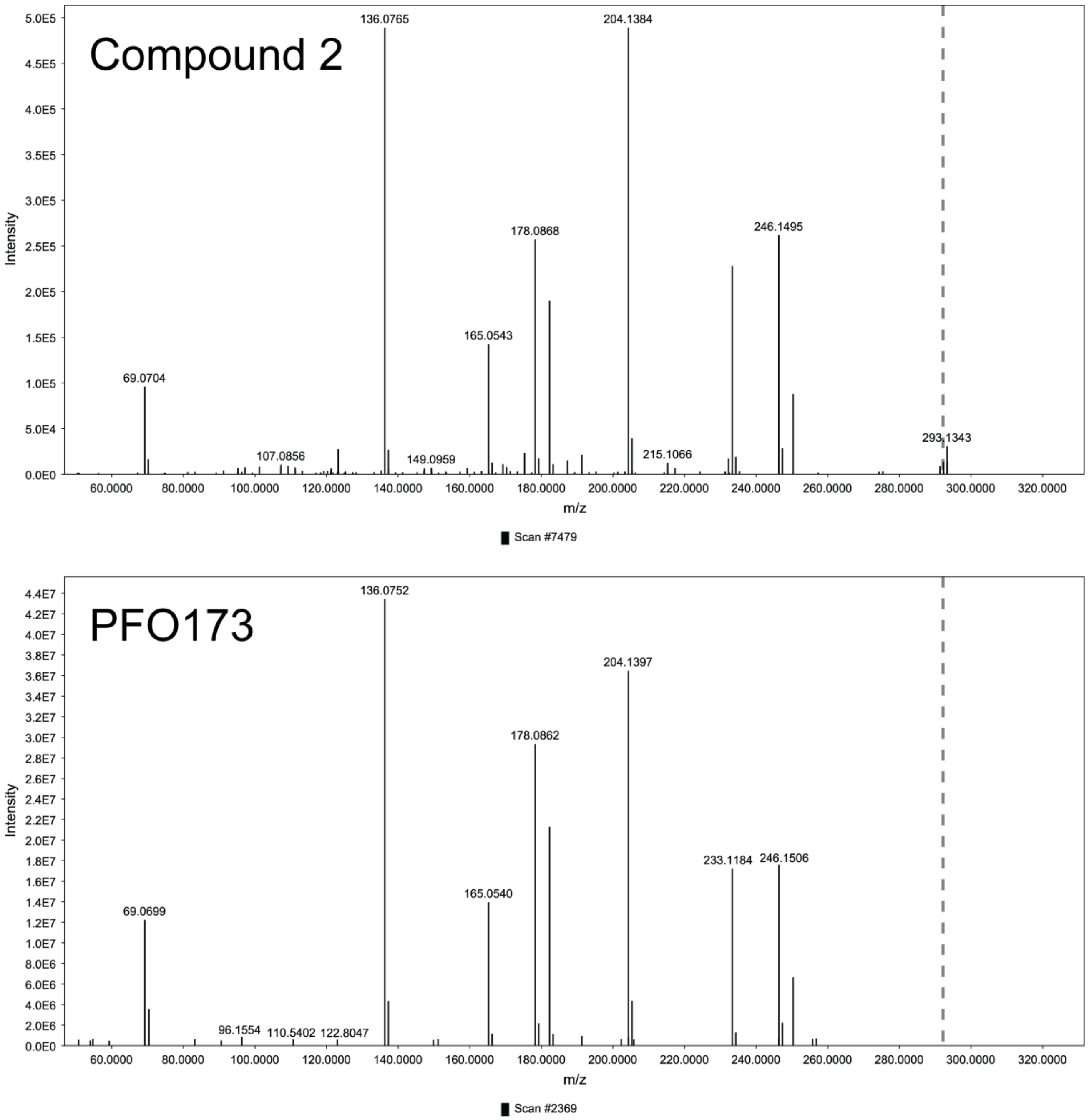


**Figure S9. Comparison of MS² spectra of compound 2 from the extract of the *Ri* DMATS overexpression strain and its chemically-synthesized reference compound, PFO173.** Precursor m/z value is indicated by a dashed grey line.

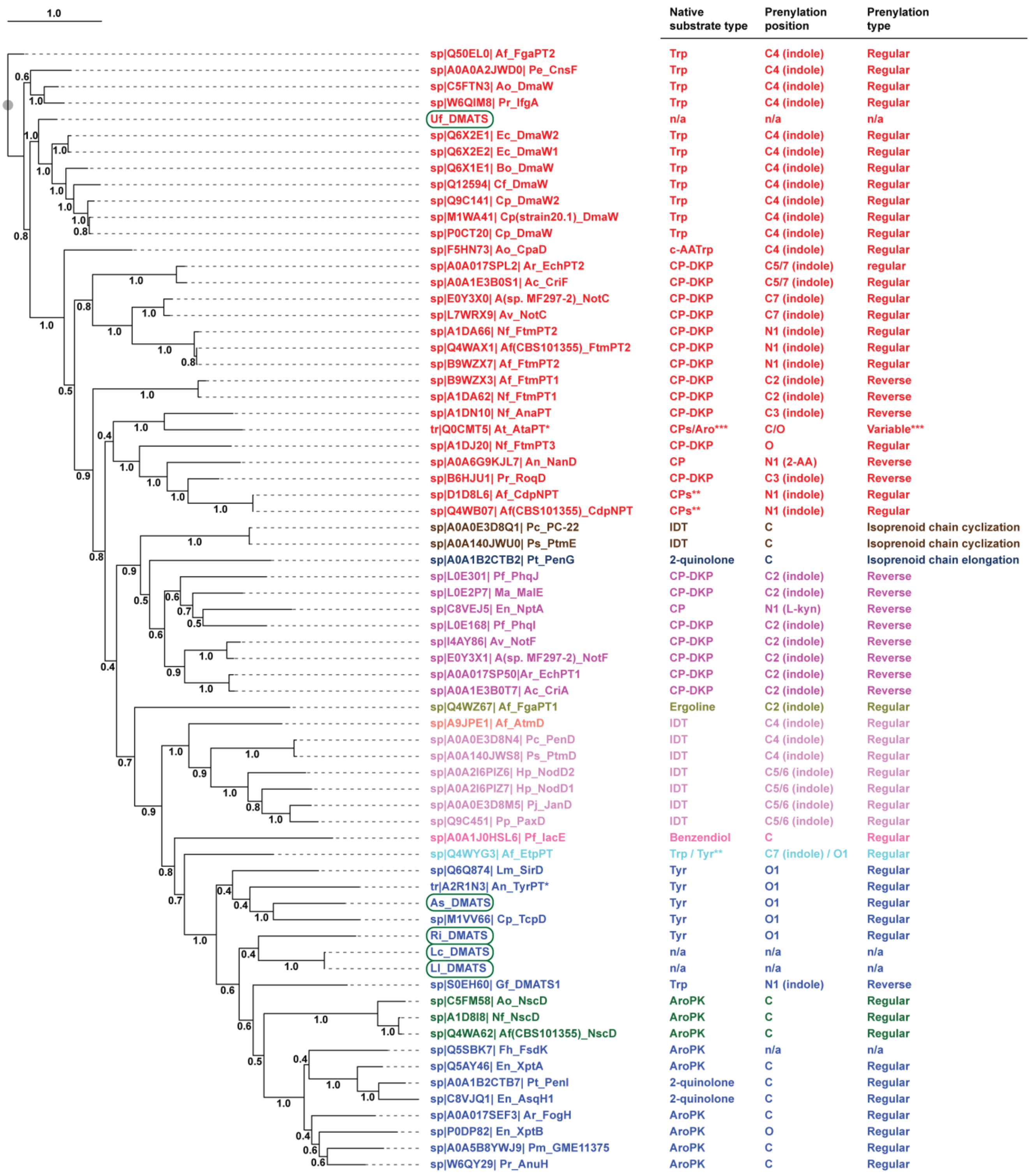


**Figure S10. Maximum-likelihood phylogenetic tree of characterized DMATS-type PTs and lichen DMATS (green circles).** *Uf* DMATS is included as the only lichen DMATS-type PT that clustered with prototype DMATS sequences. The tree was arbitrarily rooted in the *Af* FgaPT2 node. Substrates are abbreviated as follows. Trp: tryptophan; Tyr: tyrosine; c-aaTrp: cyclo-acetoacetyl-L-tryptophan; CP-DKP: cyclic peptide-diketopiperazine; CP: cyclic peptide; IDT: indole diterpene; Aro: aromatics (general); AroPK: aromatic polyketides. Entries are colored based on their respective clusters in Fig. 3. Bootstrap values are given for each node (1000 replicates). *These enzymes are not in the SwissProt database, but they have been characterized experimentally. **Unnatural substrates tested in vitro. ****At* AtaPT was only tested *in vitro* but showed an incredible range of acceptor and donor substrates [32].

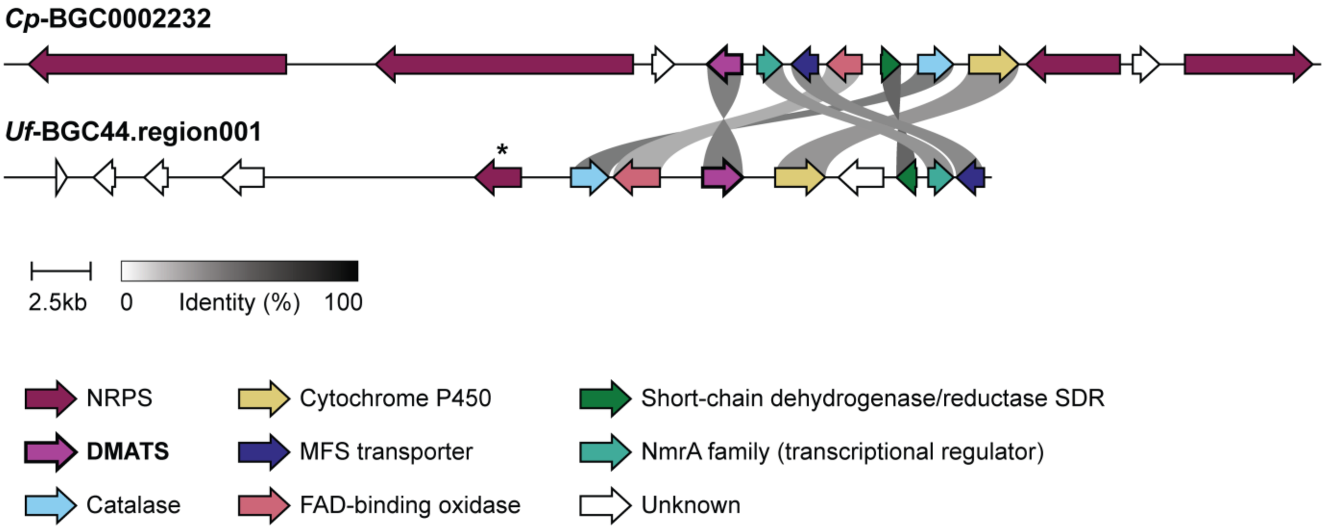


**Figure S11. Comparison of ergotamine BGC from *Claviceps purpurea* (MIBiG accession no. BGC0002232) to DMATS BGC 44.1 from *Usnea florida*.** The genes are colored based on their known or putative function. Identity between individual genes is highlighted according to provided grayscale gradient. *The putative NRPS on the BGC from *U. florida* appears to be truncated, but this might be due to inaccurate sequencing and/or genome annotation. This is also suggested by the large stretch (~9 kb) of unannotated DNA immediately after the annotated gene.

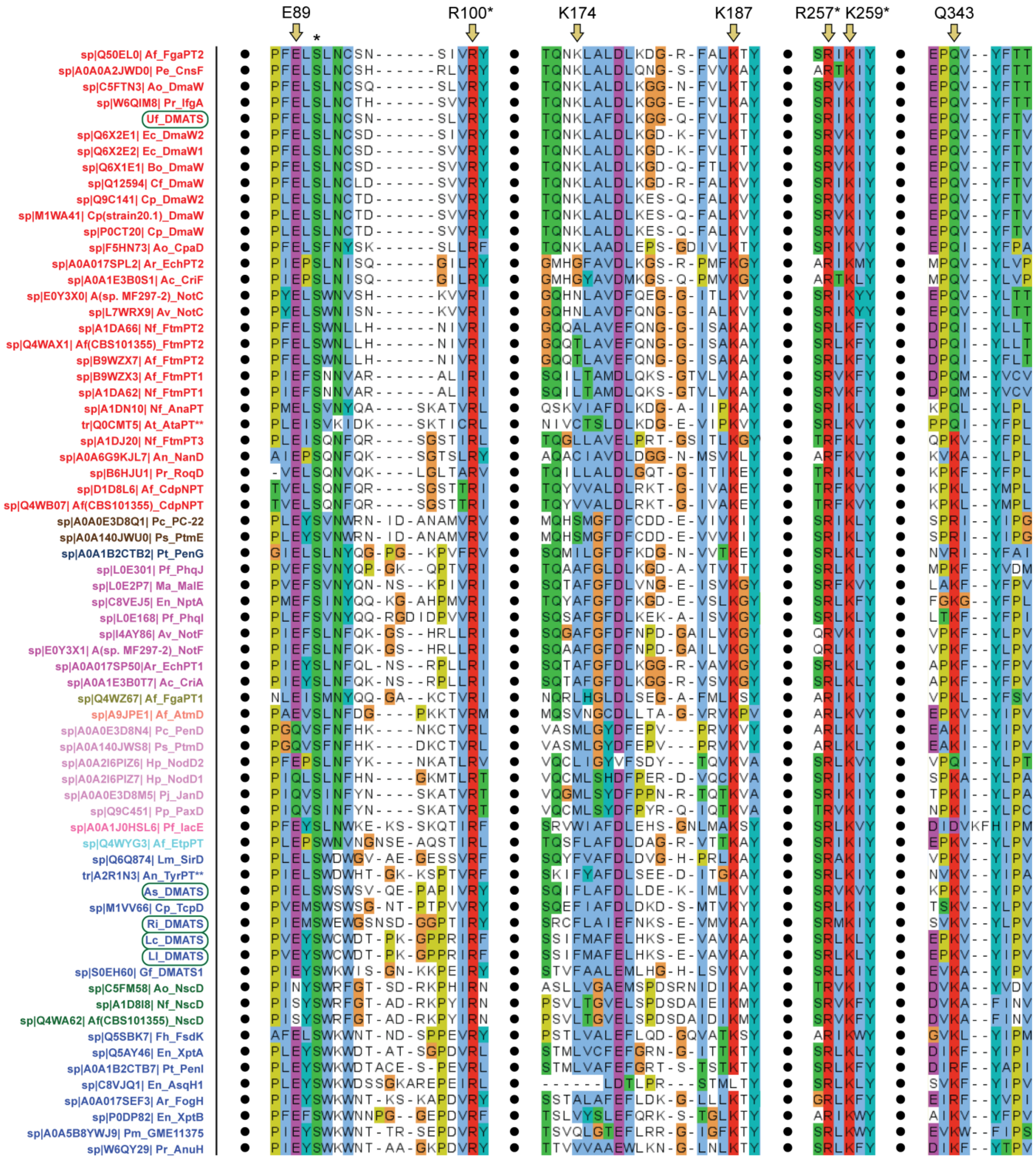


**Figure S12. Multiple sequence alignment of characterized DMATS-type PTs and lichen DMATS (green circles).** *Uf* DMATS is included as the only lichen DMATS-type PT that clustered with prototype DMATS sequences. The alignment is trimmed at relevant positions where key catalytic residues are found; the full alignment is available as additional file 2. Residues that are 100% conserved are highlighted by a star. Residues that are important for binding of substrates and catalysis are labeled and indicated by yellow arrows. **These enzymes are not in the SwissProt database, but they have been characterized experimentally.

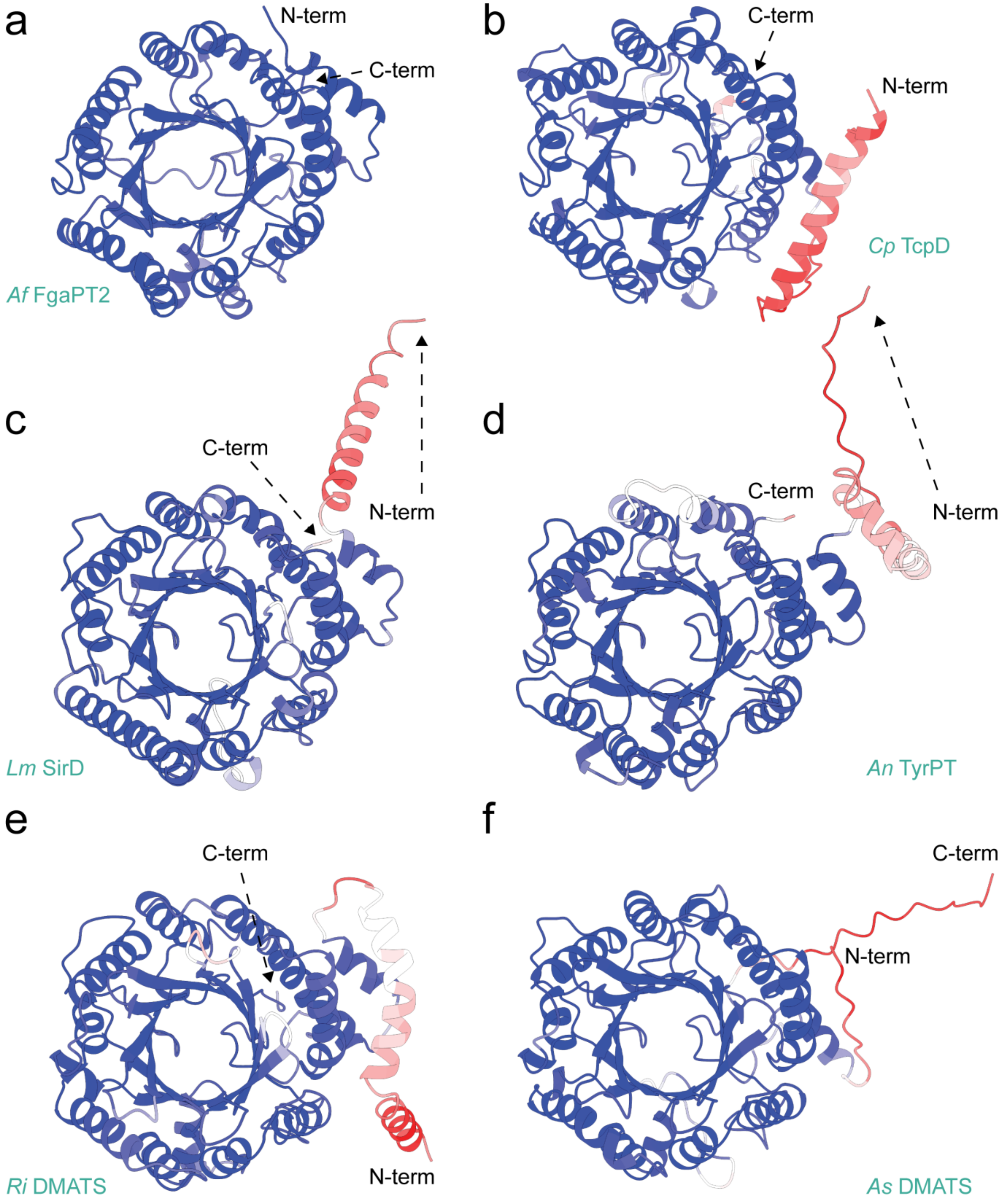


**Figure S13. Comparison of structural models of 4-O-dimethylallyltyrosine synthases (AlphaFold predictions) to the crystal structure of FgaPT2 (PDB ID: 3I4X).** The models are colored based on AlphaFold per-residue confidence measure pLDDT (higher is better, blue), while FgaPT2 is colored based on B-factor (lower is better, blue).

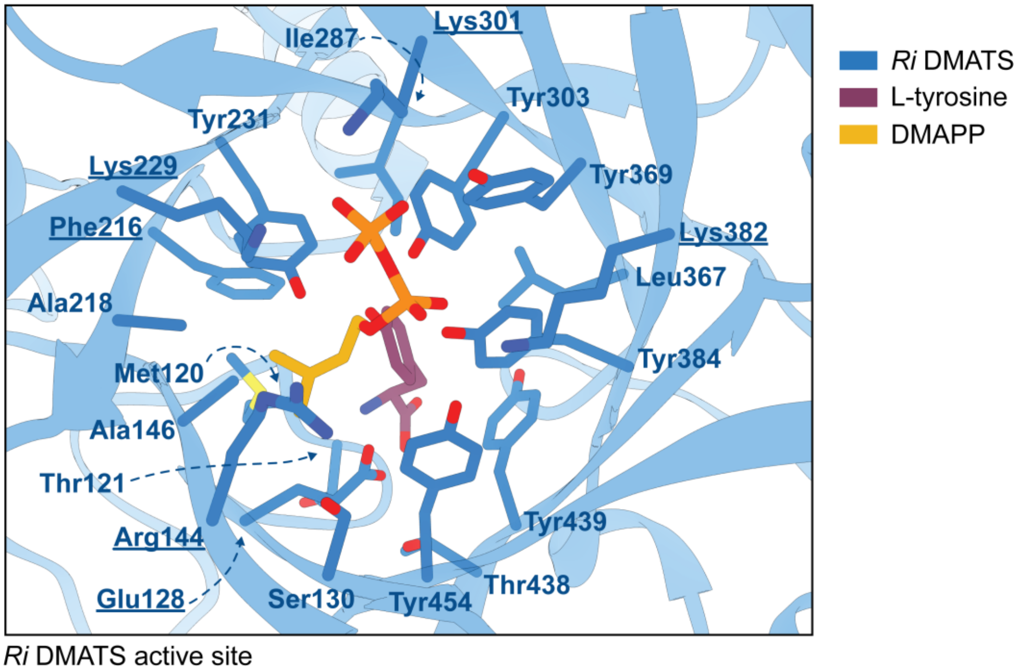


**Figure S14. Active site of *Ri* DMATS with substrates DMAPP and L-Tyr docked within the pocket.** Conserved positively charged residues Arg144, Lys229, and Lys301 interact with the pyrophosphate group as expected. The 4-OH group of tyrosine is correctly oriented to bind the dimethylallyl donor. The resulting intermediate cation could be deprotonated by a molecule of water or by Lys382, which has the ε-amino group positioned in the vicinity of the newly formed bond.
